# HDL regulates TGFß-receptor lipid raft partitioning, restoring contractile features of cholesterol-loaded vascular smooth muscle cells

**DOI:** 10.1101/2023.10.19.562786

**Authors:** Prashanth Thevkar Nagesh, Hitoo Nishi, Shruti Rawal, Tarik Zahr, Joseph M. Miano, Mary Sorci-Thomas, Hao Xu, Naveed Akbar, Robin P Choudhury, Ashish Misra, Edward A Fisher

**Affiliations:** Department of Medicine, Division of Cardiology, and Cardiovascular Research Center, NYU Grossman School of Medicine, New York, NY, United States of America; Department of Microbiology, NYU Grossman School of Medicine, New York, NY, United States of America; Vascular Biology Center, Medical College of Georgia at Augusta University, Augusta, Georgia 30912; Department of Medicine, Medical College of Wisconsin, Milwaukee, Wisconsin, USA; Division of Cardiovascular Medicine, Radcliffe Department of Medicine, University of Oxford, Oxford, United Kingdom; Oxford University Hospitals, NHS Trust, John Radcliffe Hospital, Oxford, United Kingdom; Heart Research Institute, Sydney, NSW, Australia; Faculty of Medicine and Health, The University of Sydney, NSW, Australia

## Abstract

**Background:** Cholesterol-loading of mouse aortic vascular smooth muscle cells (mVSMCs) downregulates *miR-143/145*, a master regulator of the contractile state downstream of TGFβ signaling. *In vitro,* this results in transitioning from a contractile mVSMC to a macrophage-like state. This process likely occurs *in vivo* based on studies in mouse and human atherosclerotic plaques.

**Objectives:** To test whether cholesterol-loading reduces VSMC TGFβ signaling and if cholesterol efflux will restore signaling and the contractile state *in vitro* and *in vivo*.

**Methods:** Human coronary artery (h)VSMCs were cholesterol-loaded, then treated with HDL (to promote cholesterol efflux). For *in vivo* studies, partial conditional deletion of *Tgfβr2* in lineage-traced VSMC mice was induced. Mice wild-type for VSMC *Tgfβr2* or partially deficient (*Tgfβr2+/-*) were made hypercholesterolemic to establish atherosclerosis. Mice were then treated with apoA1 (which forms HDL).

**Results:** Cholesterol-loading of hVSMCs downregulated TGFβ signaling and contractile gene expression; macrophage markers were induced. TGFβ signaling positively regulated *miR-143/145* expression, increasing *Acta2* expression and suppressing KLF4. Cholesterol-loading localized TGFβ receptors into lipid rafts, with consequent TGFβ signaling downregulation. Notably, in cholesterol-loaded hVSMCs HDL particles displaced receptors from lipid rafts and increased TGFβ signaling, resulting in enhanced *miR-145* expression and decreased KLF4-dependent macrophage features. ApoA1 infusion into *Tgfβr2+/-* mice restored *Acta2* expression and decreased macrophage-marker expression in plaque VSMCs, with evidence of increased TGFβ signaling.

**Conclusions:** Cholesterol suppresses TGFβ signaling and the contractile state in hVSMC through partitioning of TGFβ receptors into lipid rafts. These changes can be reversed by promotion of cholesterol efflux, consistent with evidence *in vivo*.

**Condensed abstract:** Many cells identified as macrophage-like in human and mouse atherosclerotic plaques are thought to be of VSMC origin. We identified cholesterol-mediated downregulation of TGFβ signaling *in vitro* in human (h)VSMCs by localization of TGFβ receptors in membrane lipid rafts, which was reversed by HDL-mediated cholesterol efflux. This restored VSMC contractile marker (*Acta2*) and suppressed macrophage marker (CD68) expression by promoting TGFβ enhancement of *miR-145* expression. *In vivo*, administration of apoA1 (which forms HDL) to atherosclerotic mice also promoted VSMC *Acta2* expression and reduced CD68 expression. Because macrophage-like VSMC are thought to have adverse properties, our studies not only show mechanistically how cholesterol causes their transition, but also suggest that efflux-competent HDL particles may have a therapeutic role by restoring a more favorable phenotypic state of VSMC in atherosclerotic plaques.

## INTRODUCTION

Atherosclerosis is a chronic inflammatory disease characterized by accumulation of lipid-laden foam cells in arteries^1, 2^. Despite advances in therapies in treating cardiovascular disease (CVD), residual risk remains with rupture of advanced atherosclerotic plaques, which leads to myocardial infarctions and strokes. Pre-clinical atherosclerosis research has predominantly focused on preventing plaque progression through reducing the number or inflammatory state of intraplaque monocyte-derived macrophages^3-5^. There has been increasing attention to vascular smooth muscle cells (VSMCs), as recent studies have extended understanding of their robust plasticity to the molecular level. Classically, it has been believed that VSMC, in addition to their contractile function in the arterial media, are also atheroprotective in the plaque intima by forming a fibrous cap to prevent rupture, in contrast to intimal macrophages, which during plaque progression, have a number of adverse effects, including foam cell formation, promotion of inflammation, and expansion of the necrotic core^6^.

The distinction between ‘protective’ and ‘detrimental’ plaque cell types, however, has blurred with the development of lineage tracing and single cell RNA sequencing techniques. As noted above, VSMC can assume multiple phenotypes ^7-9^. Current understanding is that intimal VSMCs derive from a subset of cells that clonally expand from the medial wall to assume a sub-endothelial position^10, 11^. As the plaque progresses, these protective fibrous cap cells lose their expression of typical VSMC contractile genes (such as *Acta2*, *Tagln*, *Myh11*), migrate into the intima and adopt phenotypes of various other cell types, including macrophages^12^. It is not known yet whether this plastic nature of VSMC-derived cells can be influenced to stabilize the atherosclerotic plaque and prevent rupture, or what the signals are that dynamically regulate VSMC phenotype transitions during atherogenesis.

With regard to the potential signals, the transforming growth factor beta (TGFβ) signaling pathway is of particular interest because of its well-known role in VSMC differentiation^13^. TGFβ receptor signaling is activated by the binding of TGFβ ligands to a heteromeric receptor complex composed of TGFβR1 and TGFβR2^13^. Activation of TGFβR1 leads to the phosphorylation of SMAD2 and SMAD3, which form a complex with SMAD4, which then migrates to the nucleus to influence the expression of contractile VSMC target genes, such as *Acta2*^14,15^. We were struck by the report that the conditional deletion of TGFβ signaling in VSMCs promoted phenotypic switching in an aortic aneurysm mouse model, with the appearance of cells of VSMC origin expressing macrophage markers^16^. Taken with the finding in lung epithelial cells that cholesterol treatment increased accumulation of TGFβR1 and TGFβR2 in plasma membrane domains enriched in cholesterol (i.e., lipid rafts) and decreased TGFβ signaling^17^, this suggested a potential mechanism for our previous observations that cholesterol-loading of mouse VSMC promoted the down regulation of contractile genes^18, 19^.

That cholesterol-loading may lower TGFβ signaling also in VSMC is reinforced by the findings that in loaded cells^19^ and in the aortae of hypercholesterolemic mice^20^, *miR-143/145* are downregulated. These microRNAs are positively regulated by TGFβ^21^ and are known to promote the expression of mRNAs associated with the contractile state^22^. Interestingly, *miR143/145* suppresses KLF4, a monocyte differentiation factor^22,23^. When KLF4 was knocked out in hypercholesterolemic mice, the percentage of cells of VSMC origin that expressed macrophage markers was reduced by 50%^24^. Thus, it is possible that loss of TGFβ signaling upon cholesterol loading can account for both the loss of the contractile state and the acquisition of macrophage characteristics.

If the mechanism for the suppressive effects of cholesterol loading on TGFβ signaling in VSMCs is similar to that discovered in epithelial and endothelial cells, namely the partitioning of its receptors to lipid rafts^17^, this may also provide insight into the dynamic regulation of VSMC phenotypic transitions to macrophage-like cells. Previous work has shown that high density lipoprotein (HDL)-promoted cholesterol efflux reverses the effects of cholesterol loading on mouse VSMC *in vitro^19^*. Notably, HDL reduces lipid rafts in monocytes and macrophages by depleting them of cholesterol^25, 26^. Taken together, this suggests that the reversal of the cholesterol loaded VSMC phenotype by HDL may be through its restoration of TGFβ signaling through displacement of its receptors from lipid rafts. That this may contribute to atheroprotection would be consistent with the clinical data that *functional* (i.e., efflux competent) HDL particles are associated with decreased CVD event rates (e.g., ^27, 28^) and the pre-clinical data that raising HDL particle levels promotes plaque regression and increases fibrous cap formation^29,30, 31^.

In the present study, therefore, we aimed at defining the relationships between HDL, cholesterol-loading, TGFβ signaling, *miR-143/145* expression, and VSMC phenotypes. We have extended our previous studies in mouse VSMC to human coronary artery (hVSMCs). We have also studied genetically altered atherosclerotic mice with reduced expression of *Tgfβr2*. As will be presented, cholesterol loading of hVSMCs indeed partitions the receptors into lipid rafts and impairs TGFβ signaling and *miR-143/145* expression. Furthermore, phenotypic switching of cholesterol-loaded hVSMCs to a non-contractile, macrophage-like state was reversed by increasing the levels of functional HDL particles, which displaced TGFβ receptors from lipid rafts and restored signaling. The mouse data we will present also indicate that the loss of the VSMC contractile phenotype *in vivo* may be restored when the level of functional HDL particles is raised. Taken together, our findings present TGFβ signaling as a key regulatory pathway of VSMC plasticity in hypercholesterolemic settings, with the potential to provide atheroprotection by the restoration of the signaling in intimal cells of VSMC origin.

## MATERIALS AND METHODS

### Cell Culture

Human coronary artery smooth muscle cells (HCASMC) (referred to as hVSMC) were purchased from Cell Applications and maintained in complete medium (Cat. #311-500) as provided by the vendor. hVSMCs were used within 8 passages for all experiments. Cells were cultured until 90% confluence in 37°C in a 5% CO_2_ incubator. For cholesterol or TGFβ1 treatment, cells were serum starved for 24h in 0.2% BSA (in basal media without serum, Cat. #310-500), and treatments including methyl-β-cyclodextrin-cholesterol mixture (5µg/ml, Sigma; hereafter referred to as cholesterol treatment), TGFβR1 inhibitor (SB431542, Sigma), and recombinant human TGFβ-1 (R&D Systems) were performed.

### Human high-density lipoprotein (HDL) isolation and apoA1 purification

Human plasma was obtained from NYU Langone Medical Center blood bank. HDL was isolated from plasma by a sequential flotation ultracentrifugation method. Briefly, 30ml plasma was overlayed with 20 ml of 1.019 g/ml potassium bromide (KBr) density solution in 70 ml polycarbonate centrifuge tubes and ultracentrifuged at 40,000 rpm for 24 h at 4^0^C to separate chylomicrons, IDL, and VLDL as upper fractions. The lower fraction containing LDL and HDL was collected, adjusted to 1.080 g/ml density with KBr and overlaid with 1.063 g/ml KBr density solution and ultracentrifuged at 40,000 rpm for 24 h at 4^0^C. The upper fraction (containing LDL) was removed, and the lower fraction (containing HDL and plasma proteins) was collected. Samples were adjusted to 1.225g/ml density with KBr and overlaid with 1.21 g/ml KBr solution and ultracentrifuged at 40,000 rpm for 24 h at 4^0^C. The upper fraction containing HDL was collected and stored in -80°C until apoA1 purification, as described in^32^.

### Mice

All experimental procedures were done in accordance with the New York University Grossman School of Medicine’s Institutional Animal Care and Use Committee (approved protocol # IA16-00519). ROSA26 mT/mG. Myh11-CreERT2 /J mice and ROSA26 mT/mG. Myh11-CreERT2; Tgfβr2fl/fl/J mice containing Myh11-CreERT2 inserted on the Y chromosome^33^ were obtained from Dr. George Tellides (Yale School of Medicine). For animal studies, all analyses were blinded whenever possible through numerical coding of samples.

Male mice of 8 weeks of age were intraperitoneally injected once with the mPCSK9D377Y gain-of-function transgene at 1.1×10^12^ viral particles/mouse (Penn Vector Core, University of Pennsylvania, PA). Two weeks post-PCSK9 injection, Cre-lox recombination was induced by injecting tamoxifen (Sigma) intraperitoneally at 1 mg/dose for 5 days. Mice were then placed on western diet (WD, containing 21% fat, 0.3% cholesterol, Dyets Inc.) for 20 weeks, ad libitum to develop advanced atherosclerotic plaques. Mice were monitored regularly and mice with a total cholesterol level less than 400 mg/dl were excluded from the study.

### apoA1/HDL-mediated atherosclerosis regression

Mice were randomly assigned to either progression (saline injection) or regression (apoA1 injection) groups. To promote plaque regression, mice were continued on western diet and apoA1 (500µg/mice) was administered subcutaneously twice a week for 2 weeks. Previous studies have shown that injected apoA1 rapidly associates with HDL particles^34^. Saline injections served as vehicle control.

### Plaque morphometrics and immunohistochemistry

*In vivo samples:* Aortic root sections were fixed with 4% paraformaldehyde for 15 minutes, permeabilized with 0.1% Triton X-100 for 30 minutes, followed by blocking with 3% BSA in PBS. Sections were stained with CD68 (Bio-Rad) overnight at 4°C overnight. Sections were then incubated with Alexa-Fluor 647 goat anti-rat IgG secondary antibody (Life Technologies) and stained with DAPI to detect nuclei. Images were acquired on Leica TCS SP5 confocal microscope. For some samples, sections were stained with Phospho-SMAD2 (Ser465, Ser467) Polyclonal Antibody (ThermoFisher Scientific) followed by staining with FITC Anti-GFP antibody (Abcam), and DAPI staining to detect nuclei. Image processing and quantification of the stained area were performed using Image-Pro Plus software (Media Cybernetics).

*In vitro samples:* hVSMCs were grown on sterile glass coverslips. After serum starvation (0.2% BSA in complete media) for 24h, cells were treated for 24h. Cells were washed with PBS twice and then fixed in 4% paraformaldehyde for 10 min. After being rinsed with PBS twice, cells were permeabilized with 0.1% TritonX-100 for 5min, followed by blocking in 4% normal goat serum in PBS. Anti-SMAD2/3 (Cell Signaling) were incubated at 1:200 dilution at 4^0^C overnight. Alexa-Flour 488-conjugated goat anti-rabbit IgG (Life Technologies) was used to detect SMAD2/3 localization. Then, Alexa-Flour 568-Phalloidin (Life Technologies) was incubated at 1:50 dilution for 30min. Coverslips were put on slides and mounted with medium containing DAPI. Images were acquired using Leica TCS SP5 confocal microscopy.

### Aortic digestion and flow cytometry

Mouse aortic arches were incubated in digestion buffer containing liberase (Cat. # 273582, Roche), hyaluronidase (Cat. #3506, Sigma), DNase I (Cat. # DN25, Sigma), and 1 mol/L CaCl2 at 37°C for 15 minutes using the GentleMacs dissociator (Miltenyi Biotech, Bergisch Gladbach, Germany). The digested tissue was passed through a 70µm cell strainer, washed with 1× cold PBS and centrifuged at 350 g for 10 minutes at 4°C. Cells were incubated with viability dye eFluor 780 (eBioscience, CA) for 30 minutes on ice, blocked with TruStain fcX (BioLegend, CA), and then stained with PE/Cy7 anti-mouse CD11b antibody (Cat. #101216; BioLegend) and BV605 anti-mouse F4/80 (Cat. #123133; BioLegend) for 30 minutes on ice. Following this, SMC lineage positive (GFP+) cells that were double positive for CD11b and F4/80 macrophage markers were sorted on a fluorescence-activated cell sorter (FACS) Aria II cytometer (BD Biosciences, NJ) equipped with a 100 μm nozzle, and were stored in TRIzol reagent (Invitrogen) for RNA isolation. **Real-Time qPCR** Total RNA was isolated from cultured hVSMC using TRIzol reagent (Invitrogen). cDNA was synthesized from total RNA using Verso cDNA Synthesis Kit (ThermoFisher Scientific) or Taqman MicroRNA Reverse Transcription Kit (Applied Biosystems) according to the manufacturer’s instructions. For real-time qPCR, specific mRNA or *miR-143/145* was amplified using Power SYBR Green PCR Master Mix (Applied Biosystems) or Taqman Universal PCR Master Mix, No AmpErase UNG (Applied Biosystems), respectively. Expression was normalized to *GAPDH* for mRNAs or *U6* for miRNAs.

### hVSMCs Extracellular Vesicles (EVs)

2 x 10^6^ hVSMCs were seeded in to tissue culture flasks and incubated with 15 mL of serum free complete medium (Cat. #311-500) for 24 hours under control or cholesterol treated (methyl-β-cyclodextrin-cholesterol mixture (5µg/ml, Sigma)) conditions using an established protocol^35, 36^. EVs were isolated from conditioned cell culture media using differential ultracentrifugation. Cell culture superntats were harvested and cleared of cellular debris by centrifugation at 1000 g for 10 minutes at 4°C. Cleared supernatants were transferred to new 15 mL tubes and stored at -80°C until processed. Samples were thawed on ice and centrifuged at 1000 g for 10 minutes at 4°C. Supernatants were transferred to 13.2 mL QuickSeal tubes (Beckman Coulter, California, United States) and were centrifuged at 120,000 g for 120 minutes at 4°C with a MLA55 fixed-angle rotor using an Optima MAX-XP ultracentrifuge (Beckman Coulter, California, United States). The pelleted hVSMCs EVs were resuspended in 100 μL PBS and washed in 13.2 mL PBS by ultracentrifugation at 120,000 g for 60 minutes at 4°C. Subsequently, pelleted hVSMCs EV were resuspended in 100μL of PBS (ThermoFisher Scientific, Massachusetts, United States) for subsequent analysis.

### Nanoparticle Tracking Analysis (NTA)

hVSMCs EV particle size distribution and concentration profiles were determined by NTA using a Zetaview device (Particle Metrix, Inning am Ammersee, Germany) as previously described (Akbar et al., 2022). The Zetaview measured the sample chamber from 11 different positions in two continuous cycles. The settings were set at sensitivity 80, frame 30 and shutter speed 100. Silica 100 nm microspheres (Polysciences Inc., Philadelphia, United States) were used to quality check the instrument performance daily. Prior to injection into the sample chamber, samples were diluted in PBS 1:1000.

### Western Blotting

Cells were washed with PBS twice, and protein was extracted in RIPA buffer containing protease inhibitor mixture (Sigma) and phosphatase inhibitor cocktail (Roche). Protein concentration was determined by Bradford method (Bio-Rad). Equal amounts of protein were fractionated by SDS-PAGE, transferred to nitrocellulose membranes (Whatman). The membrane was blocked with 5% non-fat milk or 5% BSA for 1 h, and then incubated with the indicated primary antibody overnight at 4°C. After a 1h-incubation with the appropriate secondary antibody, specific signals were detected by ECL chemiluminescent detection reagent (GE Healthcare). The signals were quantified by densitometry analysis (Image J). The primary antibodies used were as follows: ACTA2 (Sigma, Cat. #A2547); CNN1 (DAKO, Cat. #M3556); SRF (Cell signaling, Cat. #5147); p38MAPK (Santa Cruz Biotechnology, Cat.#sc-535); SMAD2/3 (Cell Signaling, Cat. #8685); TUBA (Sigma, Cat. #T-5168); and SMAD2 (Cell Signaling, Cat. #3103), phospho-SMAD2 (Cell Signaling, Cat. #3101S), phospho-p38MAPK (Cell Signaling, Cat. #9211S), SMAD4 (Cell Signaling, Cat. #9515); CD68 (AbD Serotec, Cat. #MCA1815); KLF4 (Cell Signaling, Cat. #12173); PU.1(Santa Cruz Biotechnology, Cat. #sc-352); TGFβR1 (Cell Signaling, Cat. #3712); TGFβR2 (Santa Cruz Biotechnology, Cat. #sc-400); Caveolin (BD Transduction Laboratories, Cat. #610059); CD71 (Cell Signaling, Cat. #13113); GAPDH (Ambion, Cat. #AM4300).

### Imaging and analysis

Images were acquired Leica TCS SP5 confocal microscope. Image processing, analysis and cell counting, were performed using Image J software.

### siRNA and miRNA Mimic/Inhibitor Transfections

*miR-143/145* mimics (60nM)/inhibitors(60nM) and siRNA (60nM) against human KLF4 (On-Target plus SMART pool siRNA) were purchased from Dharmacon. hVSMCs were transfected with 60nM of siRNA or miRNA mimic/inhibitor using RNAiMAX transfection reagent (Invitrogen) according to the manufacturer’s instructions. 24h post transfection, treatments were performed as indicated elsewhere.

### Cellular Cholesterol Measurement

Cellular lipids were extracted by using a hexane/isopropyl alcohol (3:2) mixture, followed by cellular protein extraction with 0.2 N NaOH as described^37^.Total cholesterol was determined by using kits from Wako. Total cellular protein content was determined using Bradford assay (Bio-Rad).

### TGFβ Assay

The TGFβ concentration was measured using mink lung epithelial cells stably transfected with an expression construct containing a truncated PAI-1 promoter fused to the firefly luciferase reporter gene as described^38^. The cells were kindly provided by Drs. D. Rifkin and J. Munger (New York University Grossman School of Medicine).

### Lipid raft isolation

Lipid rafts were fractionated as described^39^, followed by western blotting. All steps were performed on ice. Briefly, cells were washed and then scraped in base buffer (20 mM Tris-HCl, pH 7.8, 250 mM sucrose, supplemented with 1 mM CaCl_2_ and 1 mM MgCl_2_). Cells were subjected to centrifugation for 2 min at 250 g and the resulting pellet was resuspended in 1 ml of base buffer containing protease inhibitors. Cells were lysed by passage through a 22g × 3″ needle 20 times, and the lysates were centrifuged at 1,000 g for 10 min. The resulting post nuclear supernatant was collected and transferred to a separate tube. 1 ml of base buffer (+Protease inhibitor) was added to the cell pellet and passed through the needle and syringe for 20 times for lysis. The resulting lysate was centrifuged at 1,000g for 10 min, and the second post nuclear supernatant was combined with the first. An equal volume (2 ml) of 50% OptiPrep (diluted in base buffer) was added to the combined post nuclear supernatants and placed in the bottom of a 12 ml centrifuge tube. 8 ml gradient of 0% to 20% OptiPrep in base buffer was layered on top of the lysate, which was now 25% OptiPrep. Gradients were centrifuged for 90 min at 52,000 g using an SW-41 rotor in a Beckman ultracentrifuge. A distinct band was observed at the interface between the 20% end of the gradient and the 25% OptiPrep bottom layer. Gradients were collected into 0.67 ml fractions, and the distribution of various proteins was assessed by Western blotting.

### Statistics

All statistical analyses were performed using Prism 9 (GraphPad). *P* values were calculated using an unpaired t-test for pairwise data comparisons or one-way analysis of variance (ANOVA) for data comparisons of two or more independent groups. A p-value of ≤0.05 was considered significant.

## RESULTS

### Cholesterol loading of human VSMCs leads to downregulation of contractile gene expression

Previously we demonstrated that cholesterol loading in mouse VSMC (mVSMC) downregulated contractile gene expression^18, 19^. To extend the observations to human VSMCs, we used human coronary artery vascular smooth muscle cells (hVSMC) as a model system. Cholesterol-cyclodextrin complex was used to deliver cholesterol to hVSMC as previously^18, 19, 40^. Increases in the cellular contents of total cholesterol (Figure S1.A) and neutral lipid (presumably cholesteryl ester; CE) (Figure S1.B) confirmed the effectiveness of the loading protocol. MTT assays were performed to assess cell viability, which showed no adverse effects of cholesterol loading for at least 60h (Figure S1.C).

Consistent with our studies in mVSMC, there was a time-dependent decrease in the expression of the contractile gene smooth muscle cell marker *Acta2*, by ∼50% after 24h and ∼75% after 48h of cholesterol loading (Figure 1A). We also determined the expression for other VSMC contractile-state markers, namely Transgelin (*Tagln*) and Calponin (*Cnn1*), and these were also downregulated (Figure 1B). We have reported that in mVSMC, cholesterol loading downregulated the expression of two key transcription factors, myocardin (*Myocd*) and serum response factor (*Srf*), which govern the contractile VSMC phenotype^19^. Indeed, *Myocd* and *Srf* mRNAs were also downregulated in cholesterol-loaded hVSMCs (Figure 1B). We also confirmed the downregulation of α-SMA and CNN1 at the protein level (Figure 1C-E).

**Figure 1.**
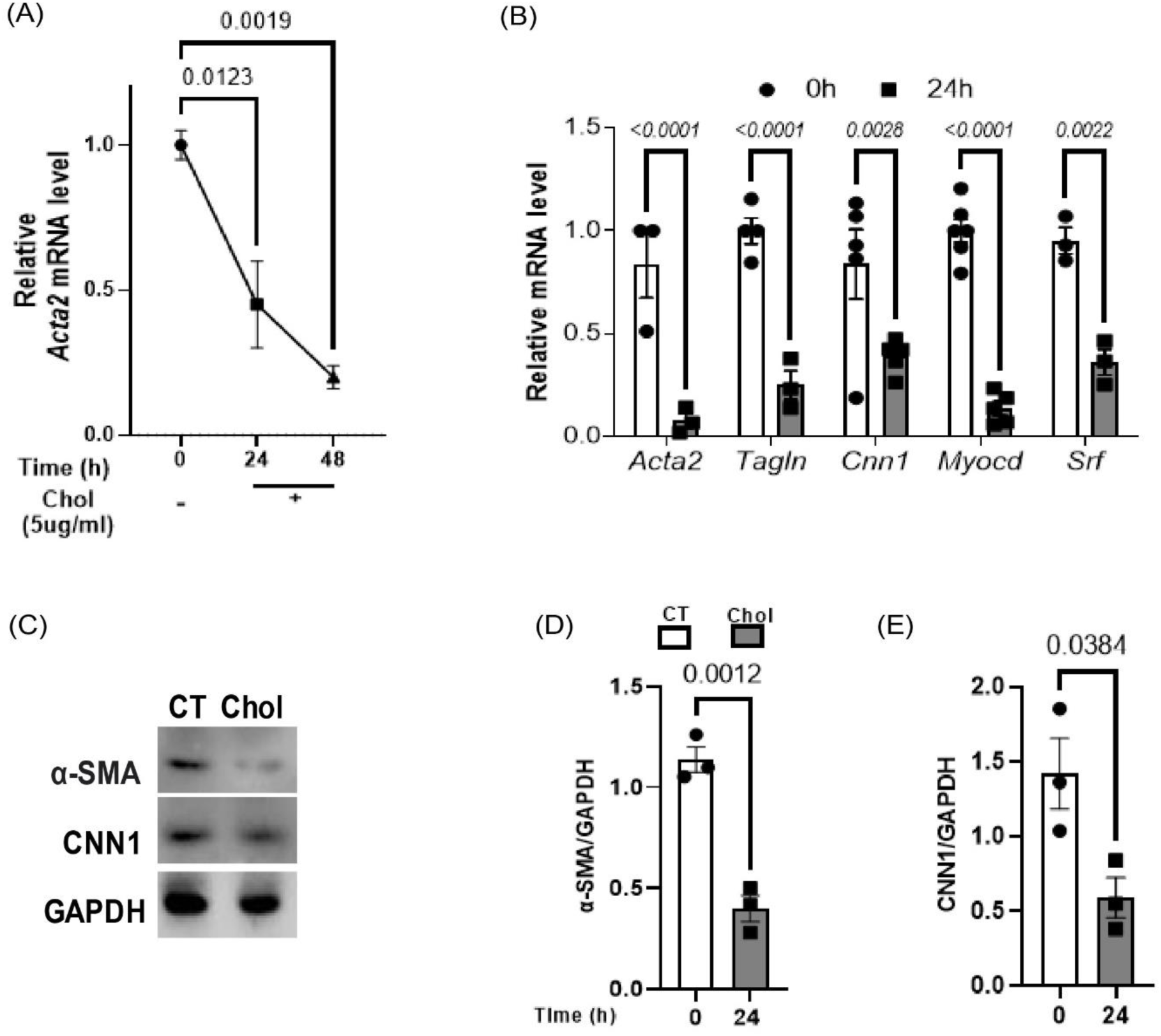
Contractile gene expression is downregulated in cholesterol-loaded hVSMC. (A-B) hVSMCs were treated with cholesterol (Chol) (5µg/ml) or 0.2% BSA (CT) for 24h and 48h and gene expression of *Acta2*, *Tagln*, *Cnn1, Myocd,* and *Srf* were determined by qPCR. (C) hVSMC were treated with cholesterol (Chol) (5µg/ml) or 0.2% BSA (CT) for 24h and protein expression of α-SMA and CNN1 were determined by Western blotting (representative blots shown). Densitometry showing the (D) α-SMA and (E) CNN1 band intensities normalized to GAPDH. Data are presented as the mean ± S.E. of three independent experiments and *p* values are as indicated.

Taken together, our data indicate that cholesterol-loading downregulates the expression of contractile-state associated genes in hVSMCs *in vitro*.

### TGFβ signaling is downregulated in cholesterol-loaded hVSMCs

Having established an attenuated hVSMC contractile phenotype with cholesterol loading, we next wanted to understand the mechanism for this. We focused on the TGFβ pathway because of its prominence in promoting the contractile state of VSMC across species^13^ and a pilot study that suggested cholesterol loading reduces TGFβ signaling in mVSMC^19^.

Additionally, we were interested in the connections between TGFβ signaling, miR143/145, and cholesterol-loading based on multiple lines of reasoning: 1) TGFβ signaling positively regulates hVSMC contractile phenotype in part by promoting the expression of miR143/145^22^; 2) Cholesterol loading in mVSMC downregulates *miR-143/145*, resulting in the loss of the contractile phenotype, and *miR-143/145* mimics protects against this loss^19^; 3) Consistent with this, *miR-143/145* expression is downregulated in aortic VSMCs in hypercholesterolemic *apoE-/-* mice^41,20^; and, in epithelial and endothelial cells cholesterol loading attenuates TGFβ signaling^17, 42^. The following series of experiments were performed, then, to test the model that cholesterol-loading reduces TGFβ signaling, which in turn decreases miR143/145 expression, resulting in the diminution of the contractile phenotype of hVSMCs.

As shown in Figure 2A and B, hVSMCs treated with TGFβ1 exhibited upregulation of the transcripts of the precursors of miR143 and 145, namely pri-miR143 and 145; strikingly, in the cholesterol-loaded cells, the responses to TGFβ1 treatment were attenuated. Furthermore, the expressions of contractile genes *Acta2* and *Tagln* were induced by TGFβ1 in control hVSMC, but this was also attenuated in cholesterol-loaded cells (Figure 2C & D).

**Figure 2.**
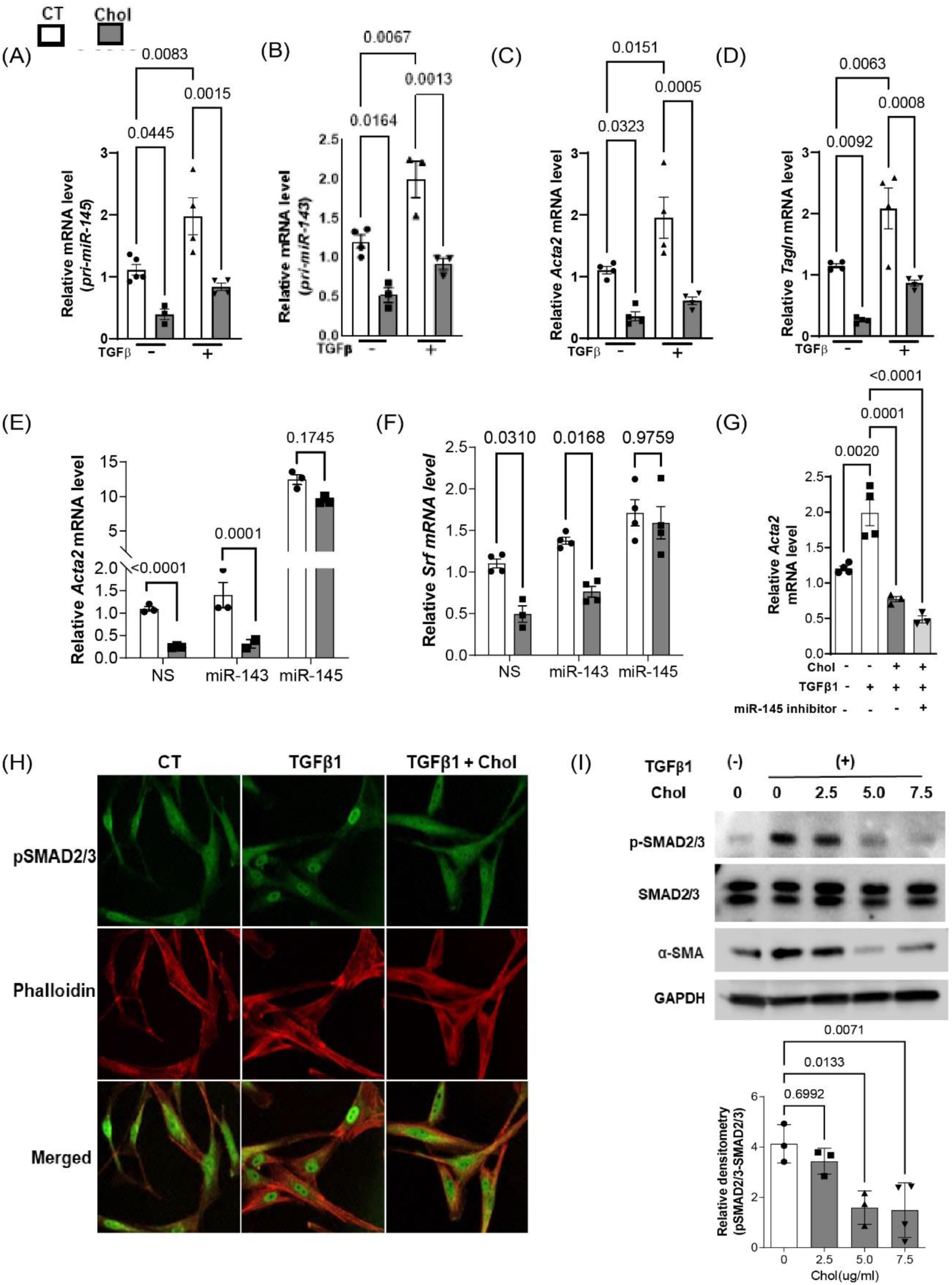
Cholesterol-loading downregulates TGFβ signaling in hVSMC. hVSMC were treated with cholesterol (Chol) (5µg/ml) or 0.2% BSA (CT; i.e., 0µg/ml cholesterol) for 24h in the presence or absence of TGFβ1 ligand (10pg/ml). Total RNA was isolated and qPCR was performed to determine the *pri-miR143/145* transcripts (A&B) or smooth muscle cell markers, *Acta2* and *Tagln* (C&D). hVSMCs were treated as in A&B, but either in the presence or absence of TGFβ1 10pg/ml) and/or non-scrambled (NS) or *miR145* mimic (60nM). qPCR was performed to determine expression of *Acta2* (E) and (F) *Srf mRNA.* (G) hVSMCs were treated as in A&B, but either in the presence or in absence of TGFβ1 (10pg/ml) and/or *miR145* inhibitor (60nM). qPCR was performed to determine expression of *Acta2.* (H) Immunofluorescence images of total SMAD2/3 (Green) in hVSMC after 24 h of the indicated treatments. Cytoplasm was stained with phalloidin (Red). Nuclei were determined as phalloidin negative area (scale bar=50µm). (I) hVSMCs were treated as in A&B, but with varying amounts of cholesterol and in the presence or absence of recombinant TGFβ1 (10pg/ml) for 24h. Proteins were extracted for western blotting to detect phosphorylated SMAD2/3 (p-SMAD2/3), and α-SMA. Total SMAD2/3 or GAPDH was used as loading controls. Blots are representative of at least three independent experiments, and the replicates were quantified by densitometry. Data are presented as the mean ± S.E. of three independent experiments and *p* values are as indicated.

Next, we over-expressed miR143 or miR145 mimics in control or in cholesterol-loaded hVSMCs. Of the 2 mimics, only miR145 upregulated the level of *Acta2* mRNA in cholesterol-loaded hVSMC to a comparable level to that in unloaded cells (Figure 2E). In addition, the miR-145 mimic prevented the suppression of *Srf,* a master regulator of the VSMC contractile phenotype in cholesterol-loaded cells (Figure 2F). Henceforth, we focused on miR145 for further studies. As shown in Figure 2G, the level of *Acta2* mRNA in TGFβ1-treated cholesterol-loaded cells was attenuated compared to that in TGFβ1-treated non-loaded cells. Notably, in the presence of a miR145 inhibitor, in cholesterol-loaded VSMCs TGFβ1 treatment failed to increase *Acta2* mRNA over that in the control cells. These results suggest that the efficacy of TGFβ1 to oppose the effects of cholesterol-loading on contractile gene expression depends on its ability to induce miR-145 (Figure 2G).

To extend these findings, we next directly determined whether cholesterol-loading decreased TGFβ1 signaling. Despite no changes in the expression levels of key downstream factors total SMAD2/3 by either cholesterol loading (Figure S2A) or TGFβ1-signaling inhibition using SB431542^43^ (Figure S2B), both of these treatments decreased contractile gene expression (*Acta2*) in hVSMCs (Figure S2C and Figure 1, respectively), and by confocal microscopy cholesterol decreased the level of active (nuclear) SMAD phosphorylated species even in the presence of TGFβ (Figure 2H). In an independent analysis (Figure 2I), the cholesterol loading-associated decrease in TGFβ1 stimulation of SMAD2/3 phosphorylation was dose-dependent, and also resulted in reduced expression of α-SMA at the protein level.

In the above experiments, exogenous TGFβ1 was added. VSMCs are known to secrete TGFβ1. To see if there is the potential for an autocrine/paracrine pathway based on endogenous production in the *in vitro* model, we measured the concentrations of both the active and latent forms of TGFβ1 in the conditioned medium of hVSMCs. As shown in Figure S3, the level of the active form (∼30 pg/mL; Figure S3A) was sufficient to activate signaling in a reporter cell assay (Figure S3B), and was in the range of the concentration of recombinant TGFβ1 that promotes SMAD2/3 phosphorylation and α-SMA induction (Figure S3C).

Overall, the results in this section support the model proposed above, namely that cholesterol-loading reduces TGFβ signaling, which in turn decreases miR143/145 expression, resulting in the attenuation of the contractile phenotype in hVSMCs. Furthermore, the pool of TGFβ1 whose signaling is being regulated by cholesterol-loading may be a component of an autocrine/paracrine process.

### Cholesterol loading partitions TGFβR1/R2 to lipid rafts and is associated with loss of TGFβ signaling

Lipid rafts are small free cholesterol (FC)-enriched portions on plasma membranes. The signaling activity of receptors can vary depending on their presence or absence in lipid rafts^44^. It has been reported that FC-loading increases lipid raft domains in multiple cell types, including VSMC^45^. Previous studies alluded to above in (mink lung) epithelial cells and (bovine aortic) endothelial cells (BAECs) have shown that when the receptor complex for TGFβ1, TGFβR1/R2 heterodimers, is in lipid rafts, TGFβ signaling is suppressed^17, 46-49^. Given the results in the previous section, implicating cholesterol enrichment of hVSMC with loss of TGFβ signaling, we hypothesized that this was because of consequent enrichment in lipid raft partitioning of TGFβ receptors.

To test this hypothesis, we determined the effects of cholesterol-loading on the distribution of TGFβR1/R2 in plasma membranes of hVSMCs. As shown in Figure 3, cholesterol loading resulted in enrichment of the receptor complex in the lipid raft region (gradient fractions 3-5), whereas in the control cells, the receptors were more abundant in the non-raft region (fractions 8-10) (Figure 3A&B). Caveolin-1 (CAV1) and the transferrin receptor (CD71) served as markers for lipid rafts, and non-raft fractions, respectively (Figure 3A)^39^.

**Figure 3.**
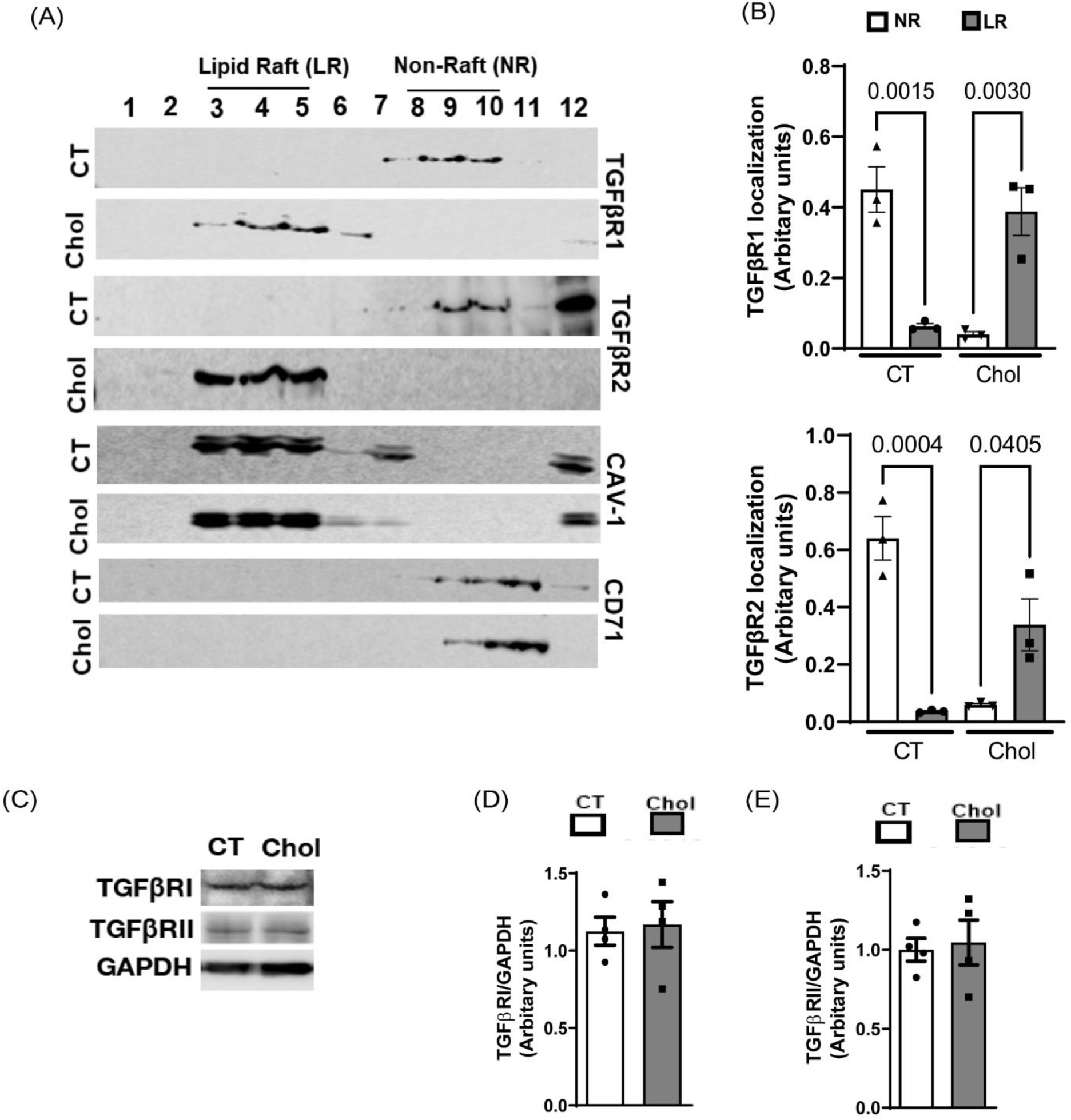
Cholesterol-loading partitions TGFβ receptors into membrane lipid rafts. hVSMC were treated with cholesterol (Chol) (5µg/ml) or 0.2% BSA for 24h. (A) Membrane lipid rafts (LR) and non-raft (NR) fractions were isolated, and Western blotting was performed using each of these fractions to determine the expressions of TGFβR1 and TGFβR2, as well as caveolin-1 (CAV1) and transferrin receptor (CD71). (B) Densitometry was performed to quantify the levels of TGFβR1 and TGFβR2 in the LR and NR fractions. (C) Western blotting was performed from total cell lysates of cholesterol-treated or untreated cells, and the bands of the TGFβ receptors visualized. (D-E) Densitometry was performed to quantify the levels of TGFβR1 and TGFβR2. Blots are representative of three independent experiments. Data are presented as the mean ± S.E. of at least three independent experiments and *p* values are as indicated.

Because extracellular vesicle (EV) shedding can be greater from lipid rafts than from other plasma membrane domains^50, 51^, another contributor to decreased TGFβ signaling could be the loss of the receptors themselves. Thus, we first determined the total EVs produced by cholesterol-loaded and unloaded hVSMCs. Indeed, cholesterol-loading increased the total number and concentration of EV-like particles released into the cell supernatants (Figure S4.A&B). Next, we isolated the EVs and performed ELISA to determine the contents of TGFβR1/R2. There was no difference in the recovery of either TGFβR1 or R2 in EVs from control or cholesterol-loaded hVSMC (Figure S4.C&D). Consistent with this were the levels of TGFβR1 or R2 in whole cell lysates, which showed no differences in their expression between control and cholesterol-loaded hVSMC (Figure 3C-E).

Overall, these results imply that it is the plasma membrane lipid raft distribution of the receptors, but not the receptor expression levels, that plays a key role in cholesterol-mediated dysregulation of TGFβ signaling.

### HDL restores the signaling of TGFβ receptors and redistributes them out of lipid rafts

We previously reported that HDL and apoA1 (the HDL-forming apolipoprotein) reversed the reduction in mVSMC contractile gene expression^19^. Therefore, we wondered if HDL restored contractile gene expression in cholesterol-loaded hVSMCs, and if so, whether it was by re-establishing TGFβ signaling by redistribution from lipid rafts. This would be consistent with the known ability of HDL to reorganize lipid rafts and modulate other signaling pathways^26, 52^.

To begin to address this, we loaded hVSMCs with cholesterol for 24h and then treated for 24 h with HDL particles (isolated from human plasma; Materials and Methods) to promote cholesterol efflux and lipid raft re-organization. As shown in Figure 4A, in hVSMC that had been cholesterol-loaded, HDL restored SMAD2 phosphorylation in response to TGFβ1 to the level observed in non-loaded cells. Furthermore, the expressions of *Myocd,* miR143/145 (Figure 4B), *Acta2* (Figure 4C), and *Cnn1* (Figure 4D) were also restored by HDL treatment in cholesterol-loaded cells. That this was related to HDL-mediated cholesterol efflux was supported by the increase in the expression of the SREBP-regulated gene *HMG-CoA reductase* (*Hmgcr*), which was suppressed by cholesterol-loading (Figure 4E). To confirm that HDL mediated restoration of the contractile pattern of gene expression via TGFβ signaling, as suggested by the SMAD2 phosphorylation results, we employed SB431542, a TGFβR1 kinase inhibitor (TGFβR1i) that decreases SMAD phosphorylation and TGFβ signaling^43^. As shown in Figure 4F, SB431542 (labeled as TGFβR1i) diminished the HDL-mediated effect on hVSMC *Acta2* gene expression.

**Figure 4.**
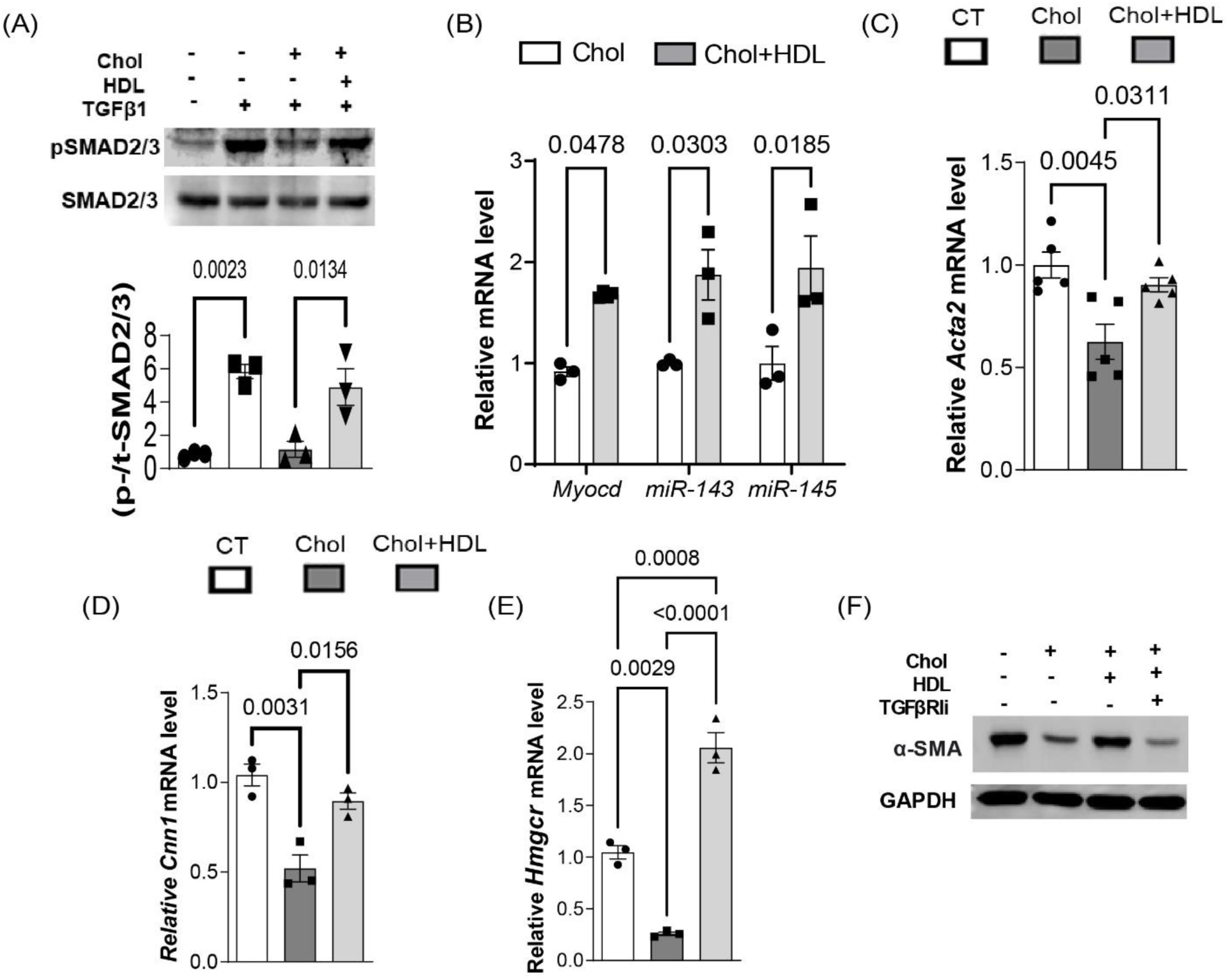
HDL treatment *in vitro* restores TGFβ signaling in cholesterol-loaded hVSMCs. (A) hVSMC were treated with cholesterol (Chol) (5µg/ml) or 0.2% BSA for 24h, followed by HDL (50µg/ml) treatment for 48h. Then, treatment groups were stimulated with recombinant TGFβ1 (10pg/ml). Western blotting was performed to detect pSMAD2 and total SMAD2, with densitometry used for quantification. (B-E) qPCR was performed to detect expression of *miR143/145, Myocd, Acta2, Cnn1* and *Hmgcr* at the conclusion of the experiment in A. (F) Cholesterol-loaded cells were either treated with HDL alone, HDL + TGFβR1 antagonist (TGFβR1i; 50ng/ml), or left untreated. Western blotting was performed to detect α-SMA. GAPDH was used as loading control. Blots are representative of at three independent experiments. *p* values are as indicated.

We next determined whether the restorations of TGFβ receptor signaling and contractile gene expression were related to HDL-induced lipid raft re-organization. As shown in Figure 5A, that HDL treatment was successful in re-organizing lipid rafts was indicated by CAV1 now being found in the dense gradient fractions, consistent with studies showing that cellular cholesterol depletion re-localizes this protein from lipid rafts to Golgi/ER membranes^53^, which are found in the bottom fractions of the sucrose gradient. Concomitant with this re-distribution of CAV1, both TGFβ receptors, which were previously enriched in lipid rafts after cholesterol loading (Figure 3), were now predominantly in the non-lipid rich fractions (Figure 5A; quantified from multiple gradients in Figure 5 B&C), where they are more active in signaling^17, 46-49^. Along with the re-distribution of the lipid rafts, there was an increase in pSMAD2 levels (reflective of increased receptor signaling) in cholesterol-loaded hVSMCs incubated with HDL (Figure 5D).

**Figure 5.**
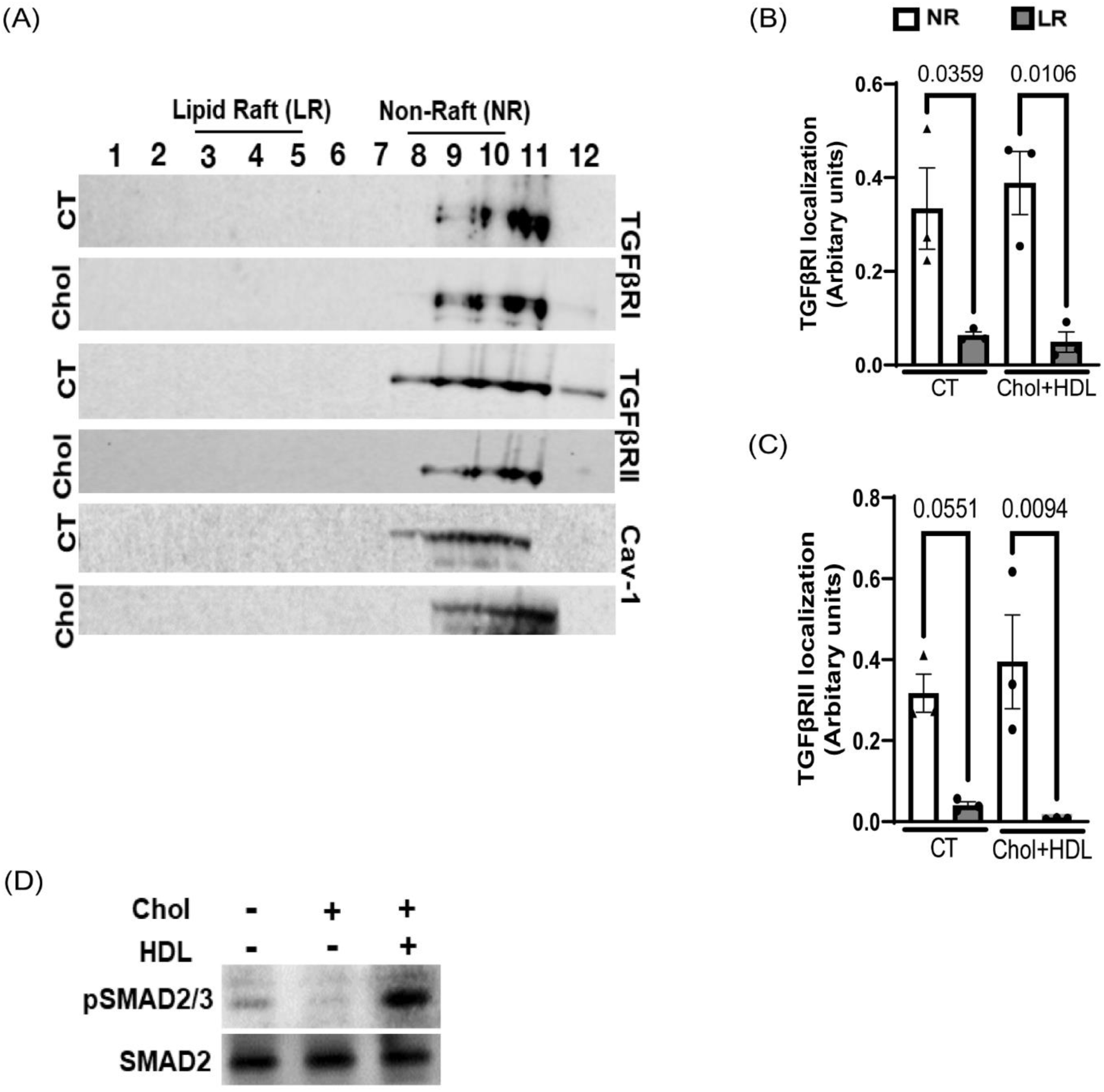
HDL treatment displaces TGFβ receptor from membrane lipid rafts in cholesterol-loaded hVSMCs and restores its signaling. hVSMC were treated with cholesterol (5µg/ml) or 0.2% BSA (CT) for 24h, after which they were either left untreated or treated with HDL (50µg/ml) for 24h. (A) At the end of the 48h protocol, lipid rafts (LR) and non-raft (NR) fractions were isolated, and Western blotting was performed using each of these fractions to determine the expressions of TGFβR1 and TGFβR2, as well as caveolin-1 (CAV1). Densitometry was performed to quantify the level of (B) TGFβR1 and (C) TGFβR2. (D) hVSMCs were loaded with cholesterol (48h, 5µg/ml), and were then either treated with HDL (50µg/ml) for 24h, or left untreated. Western blotting was performed to determine pSMAD2, SMAD2, and GAPDH levels. Data are presented as the mean ± S.E. of at least three independent experiments and the *p* values are as indicated.

### Cholesterol-loading of hVSMC promotes a macrophage-like state, which is reversed by HDL

We have previously reported in mVSMC that cholesterol-loading resulted not only in the loss of the contractile phenotype, but the assumption of a macrophage-like state^18, 19^. This was consistent with studies in mice and humans showing atherosclerotic plaques containing many macrophage-like cells (as detected by marker expression) of VSMC origin (e.g.^12, 24, 54-56^). Therefore, we sought to determine whether this could be explained by cholesterol-loading of hVSMC and, if so, what the mechanism would be.

As shown in Figure 6A, at 48h, cholesterol-loading of hVSMC again decreased the mRNA levels of *Acta2*, whereas that of *Cd68*, a commonly accepted macrophage marker, was upregulated. Note that the time courses of these changes were different, with significant decreases in the mRNA levels for *Acta2* occurring at 24h and for *Cd68* at 48h. This temporal pattern suggested that the loss of TGFβ signaling (reflected by the decrease in *Acta2* mRNA expression) likely precedes the gain in the mRNA expression of the macrophage marker *Cd68.* We hypothesized that this represented a functional link between the loss of TGFβ signaling and the gain of macrophage-like features. A prime candidate to be central in this link is KLF4, given that it is a known monocyte differentiation factor^57^ whose expression is repressed by miR143/145^22^ (which, as noted above, are induced by TGFβ signaling)^21^, and whose deficiency in mVSMC reduced the macrophage-like cells in mouse atherosclerotic plaques by ∼36%^24^.

**Figure 6.**
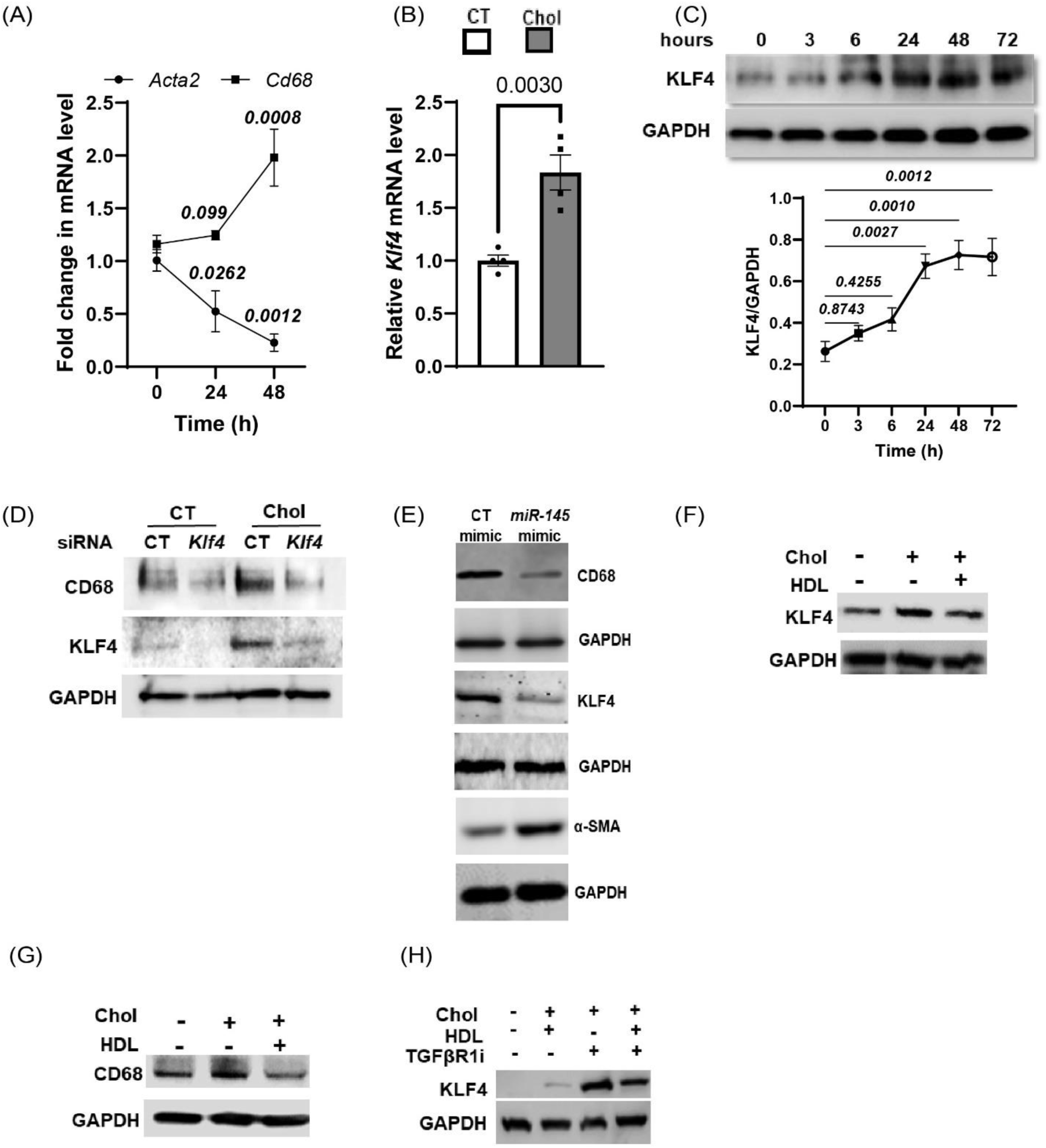
Macrophage markers upregulated in cholesterol-loaded hVSMC are suppressed by HDL through restoration of TGFβ signaling. (A) hVSMC were treated with cholesterol (5µg/ml) or 0.2% BSA (CT) for 48h. qPCR was performed to determine the expression of macrophage marker (*Cd68*) and smooth muscle cell marker (*Acta2*). (B) hVSMCs were with treated as in A for 48h, then qPCR was performed to determine the expression of macrophage differentiation factor *Klf4*. (C) hVSMCs were treated with cholesterol (5µg/ml) for the indicated times, then KFL4 expression was determined by Western blotting. (D) *Klf4* (60nM) or negative control (CT) siRNA were transfected into hVSMCs for 48h. Then, transfected cells were treated as in B, followed by western blotting for CD68 and KLF4. GAPDH was used as loading control (E) Cholesterol-loaded cells (48h, 5µg/ml) were incubated with *miR-145* mimic (60nM) or control mimic (60nM; CT) for 24h and the expressions of CD68, KLF4, and α-SMA determined with GAPDH as a loading control. (F-I) hVSMCs were loaded with cholesterol (48h, 5µg/ml), and were then either treated with HDL (50µg/ml) for 24h, or left untreated. Western blotting was performed to determine the expression of (F) KLF4, and (G) CD68. (H) hVSMCs were treated as in F&G, but in the presence or absence of TGFβR1 inhibitor (50ng/ml). Western blotting was performed to determine KLF4 expression. Data are presented as the mean ± S.E. of at least three independent experiments. *p* values are as indicated.

In an initial experiment, hVSMC were cholesterol-loaded for 48h, which resulted in upregulation of *Klf4* mRNA and protein (Figure 6B&C). Notably, the cholesterol-induced increase in CD68 was blocked by siRNA to *Klf4* (Figure 6D). We hypothesized that the effects of cholesterol loading on *Klf4* expression were a result of the reduction in miR143/145 as a consequence of reduced TGFβ signaling. This relationship was supported by the ability of a mimic of miR145 to prevent the reduction in KLF4 and CD68 in cholesterol-loaded hVSMC, while also increasing α-SMA (Figure 6E).

We next studied the effects of HDL on the phenotype of hVSMCs loaded with cholesterol. As shown in Figure 6F, KLF4 (Figure 6F) and CD68 expression (Figure 6G) were reduced by HDL. In addition, when an inhibitor of TGFβ signaling was used, the restorative effects of HDL were lost (Figure 6H).

Taken together, these results show that, as in mVSMC, cholesterol-loading promotes macrophage-like features in hVSMC, and that HDL can reverse this and restore the contractile state. Furthermore, the likely mechanism involves the restoration by HDL of TGFβ signaling, which results in upregulation of miR143/145 and repression of KLF4.

### HDL-mediated regression reduces the percentage of VSMC-derived macrophage-like cells in the advanced atherosclerotic plaque

Our results show that HDL mediates the transition of cholesterol-loaded hVSMC-derived macrophage-like cells back to a contractile VSMC phenotype by regulating TGFβ signaling *in vitro*. The current thinking is that macrophages and macrophage-like cells take up lipoproteins to form foam cells that contribute to plaque progression and inflammation^3^. There are currently no pharmacological agents that are known to drive VSMC-derived macrophage-like foam cells towards their original phenotypic state, which are assumed atheroprotective. That this issue is relevant to both pre-clinical and clinical atherosclerosis is emphasized by the reports that at least half of the foam cells with macrophage features in human plaques are VSMC-derived^54^, with similar findings in mice^55^. Thus, based on our results *in vitro*, in which cholesterol-loading of hVSMC promoted a macrophage-like phenotype and HDL reversed it, we hypothesized that a similar phenomenon could occur *in vivo*.

To test this hypothesis, we studied mVSMC lineage tracing mice (reported in^33^) with the partial conditional deletion of *Tgfβr2* (Myh11-CreERT2:ROSA26mTmG/+: TgfbrIIfl/+; hereafter referred to as *Tgfβr2*+/- mice), and induced atherosclerosis via recombinant adeno-associated virus (AAV.8) PCSK9 injection to raise cholesterol levels^58^. Mice with native TGFβ signaling (Myh11-CreERT2; ROSA26mTmG/+; *Tgfβr2*+/+), referred to as Tgfβr2+/+ mice, were used as controls. The use of partial knockdown of *Tgfβr2* in VSMCs allowed us to better determine whether HDL restored a VSMC contractile phenotype in hypercholesterolemic mice, as homozygous knockout mice manifest macrophage marker-positive cells of VSMC origin in the absence of hypercholesterolemia^16^, likely related to the total absence of VSMC TGFβ signaling.

After tamoxifen-induced recombination and AAV.8-PCSK9 injection, *Tgfβr2*+/- and *Tgfβr2*+/+ mice were fed a Western Diet (WD) for 16 weeks. One group of mice were injected with saline, which served as progression group, while another group was injected with apoA1 (500μg/dose/mice), which rapidly assembles into cholesterol-efflux promoting HDL particles^32, 34^. We found no differences in body weights between the different groups and genotypes (Figure S5A). I*n vivo* efficacy of PCSK9 injection was confimed by plasma total cholesterol (Figure S5B). An increase in plasma HDL cholesterol was observed in apoA1 injected mice, confirming the *in vivo* assembly of HDL from apoA1 (Figure S5C). Treatment was given every 2 days for 2 additional weeks of WD feeding (Figure 7A).

**Figure 7.**
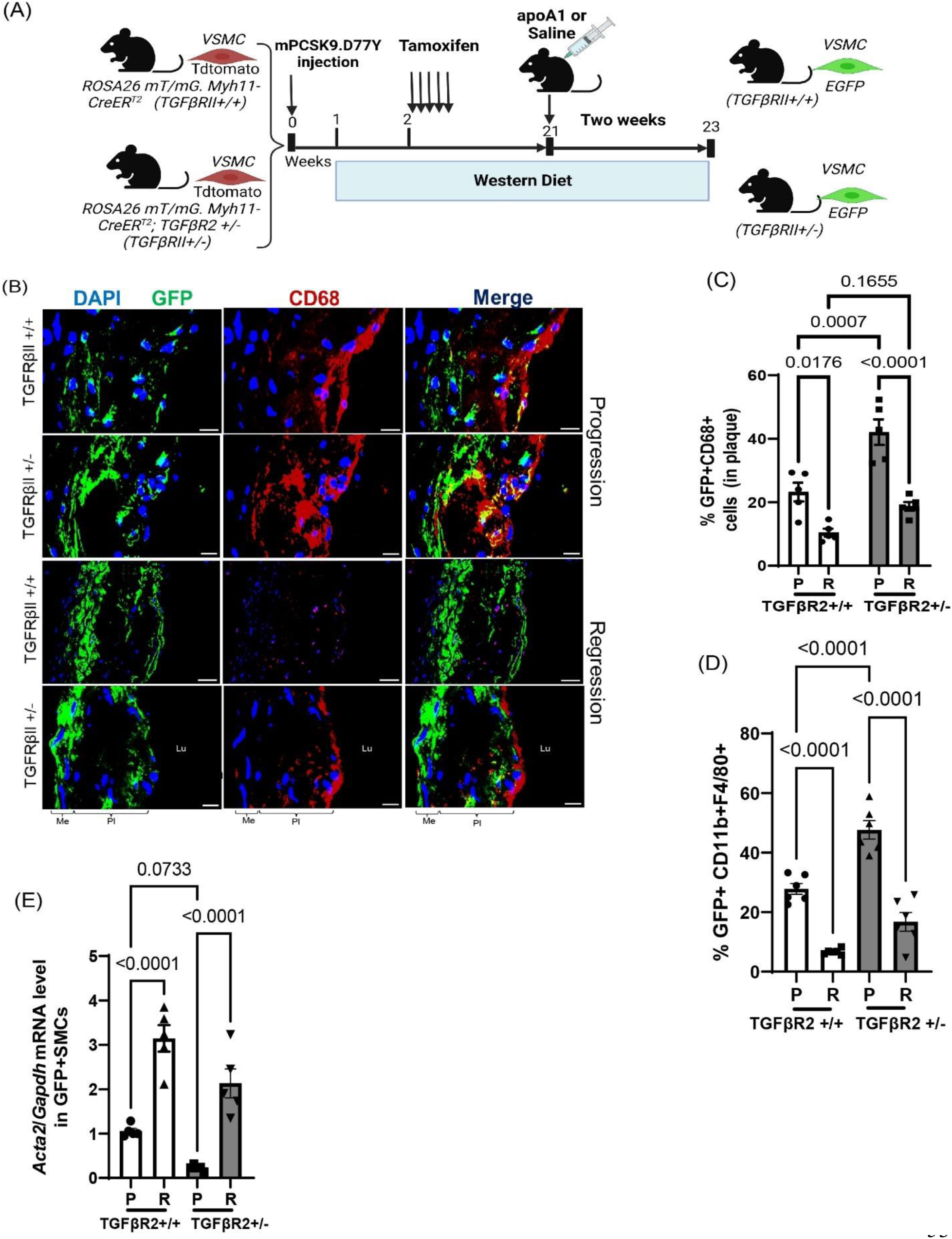
HDL increases the expression of *Acta2* relative to that of CD68 in atherosclerotic mice. (A) Schematic representation of experimental design. Note that apoA1, which forms HDL particles *in vivo,* was injected after atherosclerosis progression (P) to induce regression (R). (B) Representative images from progression (P) and regression mice (R), that were sufficient (*Tgfβr2+/+)* or haplosufficient (*Tgfβr2+/-*) for TGFR2, showing the lineage-positive VSMCs (GFP+) expressing macrophage marker or CD68 (red). Yellow color represents GFP-expressing CD68+ cells. (C) Quantification of GFP/CD68 double +. (D) Aortic digestion followed by cell sorting of GFP+ cells was performed using flow cytometry to capture lineage-positive cells (GFP) expressing macrophage markers (CD11b and F4/80). (E) Total RNA was isolated from sorted cells and qPCR was performed to identify *Acta2* gene expression. Data are presented as the mean ± S.E. (n=5-6 mice per group). *p* values are as indicated.

**Figure 8.**
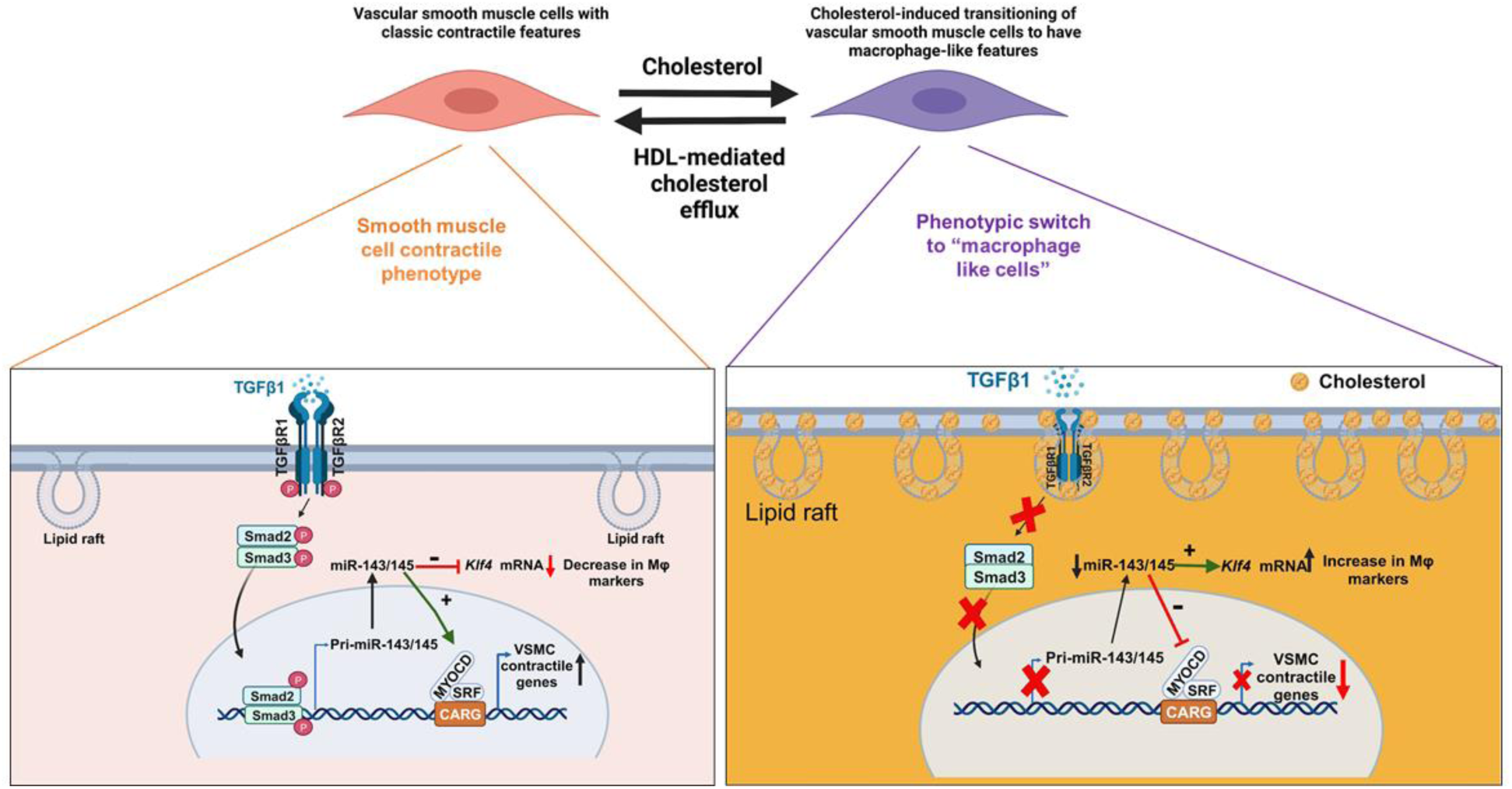
Schematic representation of proposed role of cholesterol mediated regulation of TGFβ signaling.

As expected, in the progression group, *Tgfβr2*+/- mice displayed a 20% increase in GFP+ CD68+ cells within the plaque compared to the *Tgfβr2*+/+ mice, indicative of increased mVSMC assumption of a macrophage-like phenotype (Figure 7B&C). After the injections of apoA1, the *Tgfβr2*+/+ mice and *Tgfβr2*+/- mice exhibited 10% and 22% decreases (Figure 7C), respectively, in GFP+ CD68+ cells, suggesting that the macrophage-like phenotype of the plaque mVSMCs underwent at least partial reversion to the contractile phenotype. In an independent analysis, we found that SMCs (GFP+cells) expressed higher percent of macrophage markers (GFP+CD11b+F4/80+) in *Tgfβr2*+/- as compared to *Tgfβr2*+/+ (Figure 7D). Conversely, in regressing mice, the % of GFP+CD11b+F4/80+ cells was significantly reduced in *Tgfβr2*+/+ compared to its corresponding progression group (Figure 7D). Similar changes in the % of GFP+CD11b+F4/80+ cells were found in *Tgfβr2*+/- in regression as compared to their corresponding progression groups (Figure 7D). Similarly, we found comparable increases in *Acta2* expression in both *Tgfβr2*+/+ and *Tgfβr2*+/- in mice injected with apoA1 (regression), compared to their respective progression groups (Figure 7E). That these changes were associated with increased TGFβ signaling, we also quantitated the number of cells immunopositive p-SMAD2. As shown in Fig. S6, apoA1 treatment was indeed associated with trends of increased positivity in both the mice WT and haploinsufficent for *Tgfβr2* in plaques and in the media.

## DISCUSSION

VSMCs in normal arteries have been studied typically for their contractile functions. In atherosclerotic plaques, these cells migrate from medial layer into the intima and then proliferate^10, 59^, where they can assume many fates. For example, these cells are major contributors to the smooth muscle actin+ (SMA+), collagen secreting cell population, thereby governing fibrous cap thickness and plaque stability^60-62^. Another fate in both mouse and human plaques is the acquisition of macrophage and macrophage foam cell-like features, either directly *in vitro* or after a transition *in vivo* to a multipotent SEM (“stem, endothelial, and monocyte”) cell (reviewed in^59^). Cells of VSMC origin are estimated to be a significant proportion (as high as ∼60-70%) of the macrophage marker+ cell population in plaques^55^. While the functional consequence of this is still a topic of speculation, the sheer abundance of these cells has called attention to the process whereby they originate, whether the process is reversible, and whether they contribute to the risk of adverse clinical events.

The present study provides insights into the mechanisms of hVSMC loss of the contractile phenotype and the transition to the macrophage-like phenotype in atherosclerotic plaques. First, we show that cholesterol-loading of hVSMCs dampens contractile and enhances macrophage gene expression in a time-dependent manner by impairing TGFβ signaling. Furthermore, an important consequence of this impairment is the decrease in the expression of miR143/145, important molecular factors for the positive maintenance of the VSMC contractile phenotype and suppression of the macrophage-like features. The suppression of TGFβ signaling by cholesterol-loading was driven by TGFβR1/2 enrichment in lipid rafts. This result is supported by evidence that epithelial and endothelial cells that are cholesterol-loaded also undergo TGFβR localization to rafts and suppression of TGFβ signaling (reviewed in^17^). It should be noted that another “negative feedback” regulator of TGFβ signaling in VSMCs has recently been described^63^, in which the protein LM07, initially induced by TGFβ after vascular injury, subsequently reduces TGFβ transcription. Thus, depending on the context, vascular injury or hypercholesterolemia, these and other mechanisms to reduce TGFβ signaling may be operative^64^.

We also demonstrate that treatment of cholesterol-loaded hVSMCs with HDL re-partitions TGFβRs to non-raft membrane domains, resulting in increased TGFβ-induced downstream signaling and culminating in increased miR143/145 expression. This restored the contractile phenotype and suppressed KLF4-induced macrophage marker expression. These changes were likely related to the known ability of apoA1 and HDL to deplete lipid rafts in monocytes and macrophages by promotion of cholesterol efflux^25, 26^, which is consistent with finding that HDL did not have significant effects on the expression of miR143/145 or *Klf4* in ABCA1-deficient mVSMCs^65^. To extend our findings to the *in vivo* setting, we used a murine model of atherosclerosis with VSMC lineage marking. Consistent with the data *in vitro*, TGFβR-haploinsuffiency increased the proportion in plaques of macrophage marker+ VSMC cells, and apoA1 injections, which we have previously shown to rapidly deplete plaques of cholesterol^34^, decreased this proportion and increased *Acta2* expression. Furthermore, these changes were associated with evidence that TGFβ signaling was increased in the plaques and the adjacent media. VSMC-derived macrophage-like cells have been proposed to promote plaque inflammation through a variety of mechanisms^15^ and has led to consideration of strategies to revert this phenotype. The results of our studies suggest that apoA1 or HDL particles can accomplish this, given their success to effect favorable changes in cholesterol-loaded mouse^19, 65^ or hVSMCs (this study). This would also be consistent with genetic and clinical studies that have found an atheroprotective relationship not with plasma HDL cholesterol levels, but, rather, with the cholesterol efflux function of HDL particles (reviewed in^66^). The results with the treatment of mice with apoA1 not only extend the *in vitro* results, but also suggest that the decreased expression of ABCA1 in intimal VSMCs reported in mouse and human plaques^54, 55^ does not preclude the benefits of functional HDL *in vivo*, either because the level we used overcame this deficiency, or that efflux was accomplished by one of the other well characterized routes of HDL-mediated efflux, such as through ABCG1, aqueous diffusion, or SR-B1^67^. Indeed, studies *in vivo* have shown that cholesterol flux after administration of HDL particles is preferentially mediated by SR-B1 and ABCG1^67^.

Because a key consequence of HDL treatment was its induction of *miR143/145* in cholesterol-loaded hVSMCs, a potential approach to maintaining or restoring the contractile state of VSMCs in atherosclerotic plaques could focus directly on these microRNAs rather than on factors upstream of them. Indeed, there are two recent studies in which micelles containing miR145 were used to treat either hVSMCs isolated from atherosclerotic plaques or mice with atherosclerosis^41, 68^. In the former study, the authors found that with increasing disease severity, patient-derived hVSMCs had decreasing levels of contractile markers and increasing levels of KLF4^68^. Notably, treatment with *miR145* micelles rescued contractile marker expression to baseline levels. In the mouse study, treatment with *miR145* micelles increased mVSMC contractile marker expression, collagen content and reduced necrotic core in both early atherosclerosis progression and in well-established disease^41^. One caveat is that the micelles target CCR2, which, in addition to macrophage-like mVSMCs, would also be expected to result in uptake by monocytes and macrophages.

These results taken with the favorable effects of apoA1/HDL on TGFβ signaling *in vitro* and on *Acta2* and CD68 expression *in vivo*, strengthen the case for therapeutic approaches to boosting of TGFβ signaling if the macrophage-like state is established as deleterious. Besides the present data, such an approach is supported by the growing body of literature implicating impaired TGFβ signaling in the promotion of vascular disease. For example, deletion of *Tgfβr2* in mVSMCs in *Apoe^-/-^*-deficient mice worsened plaque burden and increased the frequency of phenotypic switching to a macrophage-like cell^69^. In normocholesterolemic mice, the deletion of *Tgfβr2* in VSMCs induced the appearance of macrophage markers in mVSMC in the mouse aortic wall during aneurysm formation^16^. Additionally, in atherogenic conditions, Smad3 deficiency in mVSMCs promoted chondrogenic and ECM-remodeling phenotypes^70^. In contrast, mVSMC-specific knockdown of the transcription factor Zeb2 increased the chromatin accessibility of TGFβ signaling mediators and beneficially modulated SMC phenotype^71^.

While the observations from the Simons lab (Chen et al.^69^), demonstrating an increase in macrophage-like VSMCs after TGFβ impairment, aligns with the present findings, it should be noted that in a report from the Quertermous lab (Cheng et al.^71^) found no evidence of a VSMC-derived macrophage-like transition in mouse atherosclerotic plaques. This could be a result of the chosen VSMC model, as we and Chen *et al*.^69^ utilized conditional *Tgfβr2* deleted mice, whereas Cheng et al.^71^ conditionally knocked out the downstream mediator Smad3. Since Smad3 may be activated by other pathways (eg., non-canonical TGFβ signaling), this could limit the overlap in the observed phenotypes. Consistent with this are the differences highlighted during murine development, wherein *Tgfbr2^-/-^* mice are embryonically lethal^72^, but *Smad3^-/-^* mice are not^73^, suggesting that signaling through these two molecules have fundamental differences.

In conclusion, cholesterol-loading promotes hVSMC lipid accumulation, which results in a loss of the contractile and gain of a macrophage-like phenotype by impairing signaling of TGFβ through the partitioning of its receptors to lipid rafts. We also show for the first time that apoA1/HDL can restore TGFβ signaling in VSMCs in high cholesterol environments not only *in vitro,* but also *in vivo.* These results, taken with the literature, collectively suggest that modulating TGFβ signaling in VSMCs within the plaque milieu may be an important target for the development of atheroprotective therapeutics.

### Study limitations

As in all studies, there are limitations. For example, while the data *in vitro* provide mechanistic evidence for how cholesterol-loading and HDL treatment regulates hVSMC phenotypes, the evidence *in vivo* is associative, and more direct data will be needed to establish the findings conclusively. More advanced genetic models would benefit such studies, such as the dual lineage approach recently developed using Myh11-Dre and Cd11b-CrexER^74^. In addition, VSMCs can convert to a variety of phenotypes besides a macrophage-like state, including osteoblast-like and fibroblast-like phenotypes within the plaque^56, 71, 75-77^. While we explored the reversion of macrophage-like VSMCs, future studies should include a variety of phenotypes.

### Conclusion

Our studies highlight the loss of TGFβ signaling and its consequences in VSMCs in a high cholesterol environment, as well as the the therapeutic potential of HDL or apoA1 to restore VSMC TGFβ signaling to beneficial effect.

### Clinical perspectives

#### Competency in medical knowledge

VSMCs exhibit remarkable phenotypic plasticity in human and mouse atherosclerosis, with estimates of over half of macrophage-appearing cells being of VSMC origin. TGFβ signaling is a major regulator of the VSMC contractile state, and in pre-clinical studies, its loss in VSMCs results in a loss of contractile features and the promotion of a macrophage-like state. It is thought that these cells have adverse effects in atherosclerotic plaques. This is the first study to demonstrate in human coronary artery VSMC that cholesterol loading of the cells, as would occur in atherosclerosis progression, downregulates TGFβ signaling by localizing its receptors into membrane lipid rafts, where they are relatively inactive. Furthermore, cholesterol efflux displaces the receptors from rafts and restores TGFβ signaling, the expression of contractile genes, and the suppression of the macrophage-like state. Similarly, infusion of apoA1 (which forms cholesterol-efflux competent HDL) into atherosclerotic mice increases the balance between contractile vs. macrophage features with evidence of increased TGFβ signaling.

#### Translational outcomes

The loss of the contractile state and the acquisition of macrophage-like features in arterial VSMCs during atherosclerosis progression is thought to have adverse effects. The present studies not only provide insights into mechanisms how cholesterol levels can regulate this process, but also highlight a novel potential benefit of functional HDL particles, namely, their ability to protect against the effects of cholesterol loading on VSMC phenotype, as evidenced in studies *in vitro* (human coronary artery VSMCs) and *in vivo* (mice with atherosclerosis). Given the continued interest in HDL-based therapies, the present results may stimulate further efforts for this approach.

### Highlights

- In human coronary artery vascular smooth muscle cells (hVSMCs) cholesterol-loading downregulates TGFβ signaling and its downstream target *miR-145,* resulting in loss of contractile state and gain in macrophage-like state.
- Cholesterol induced downregulation of TGFβ signaling is due to localization of receptors TGFβR1 and TGFβR2 into membrane lipid rafts. HDL mediated cholesterol efflux displaced the receptors from lipid rafts, restored TGFβ signaling and *miR-145* expression, which resulted in restoring the hVSMC contractile state.
- In a mouse model of atherosclerosis in which VSMC are partially deficient in TGFβR2, infusion of apoA1 (which forms HDL) increased the ratio of contractile to macrophage marker expression, with evidence of increased TGFβ signaling.

## Abbreviations and Acronyms

VSMC(s): Vascular smooth muscle cell(s)
hVSMC(s): Human vascular smooth muscle cell(s)
mVSMC(s): Mouse vascular smooth muscle cells
HDL: High density lipoprotein
TGFβ: Transforming growth factor ß
TGFβR1: Transforming growth factor **β** Receptor 1
TGFβR2: Transforming growth factor **β** Receptor 2
PSCK9: Proprotein convertase subtilisin/kexin type 9

## ACKNOWLEDGEMENTS

These studies were supported by the following grant funding: N.A. and R.P.C: British Heart Foundation (BHF) Centre of Research Excellence (RE/13/1/30181 and RE/18/3/34214); BHF Project Grant (NA and RPC: PG/18/53/33895) and a BHF Intermediate Fellowship (NA: FS/IBSRF/22/25110). E.A.F.: NIH R01HL084312. J.M.M.: NIH R01HL147476. A.M.: UK-HRI grant UKIG001; Vanguard Heart Foundation grant NHF1017. M.S.T.: NIH R01HL138907. P.T.N acknowledges his late father Sri. Nagesh Thevkar for all his motivation and sacrifice leading to P.T.N’s scientific career including this manuscript.

