## Supplementary data for "HDL regulates TGFß-receptor lipid raft partitioning, restoring contractile features of cholesterol-loaded vascular smooth muscle cells"

**Supplementary Figure 1. hVSMC loaded with cholesterol exhibited increased lipid content**

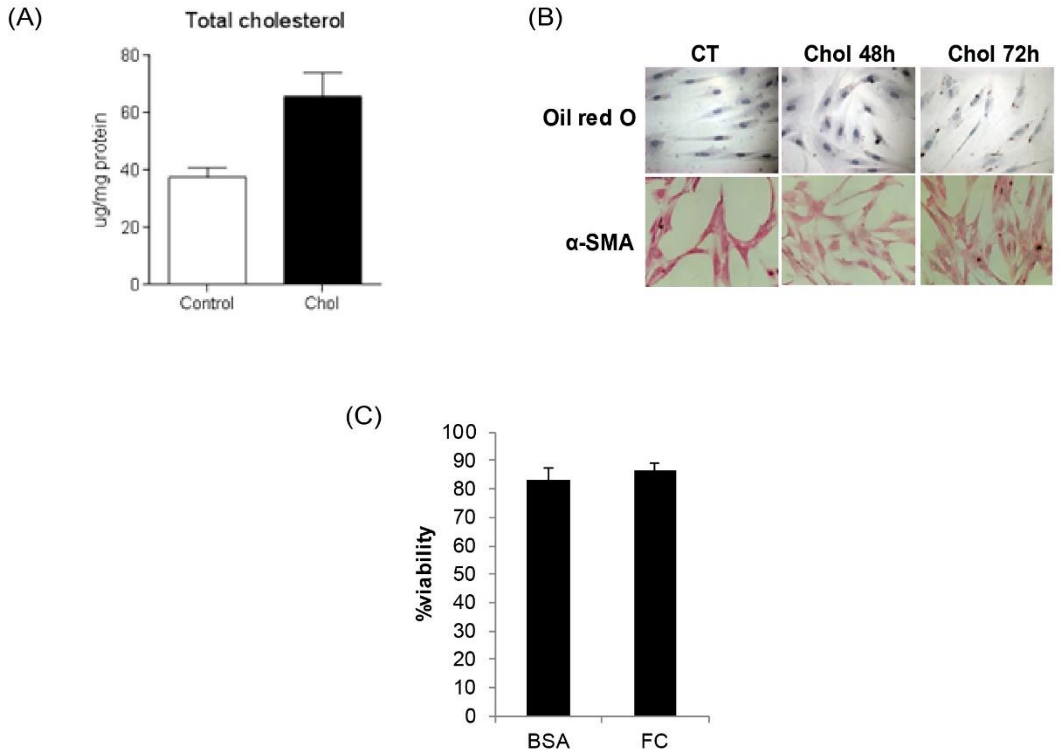

### Supplementary Figure 2. Cholesterol loading or using TGFβR1 antagonist did not affect the expression of total SMAD2/3 in hVSMCs

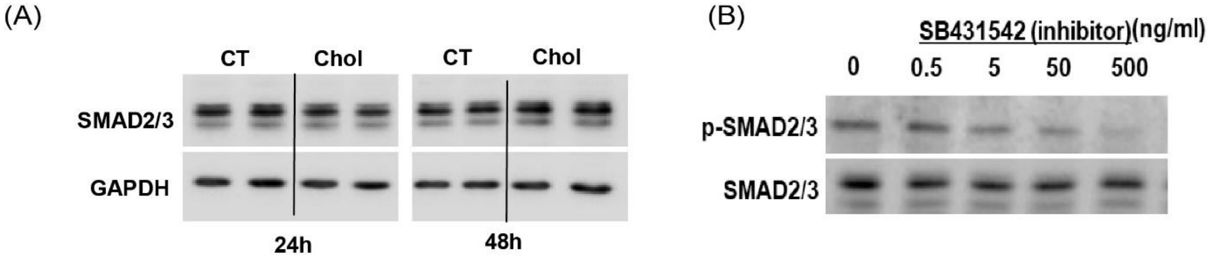


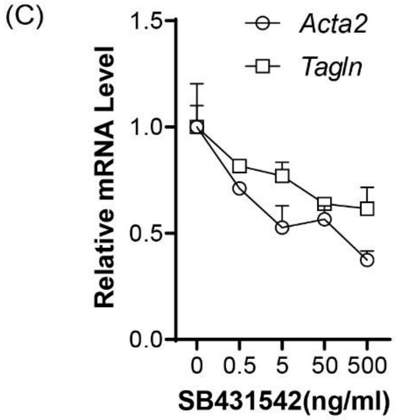

### Supplementary Figure 3. hVSMCs produce active form TGFβ1

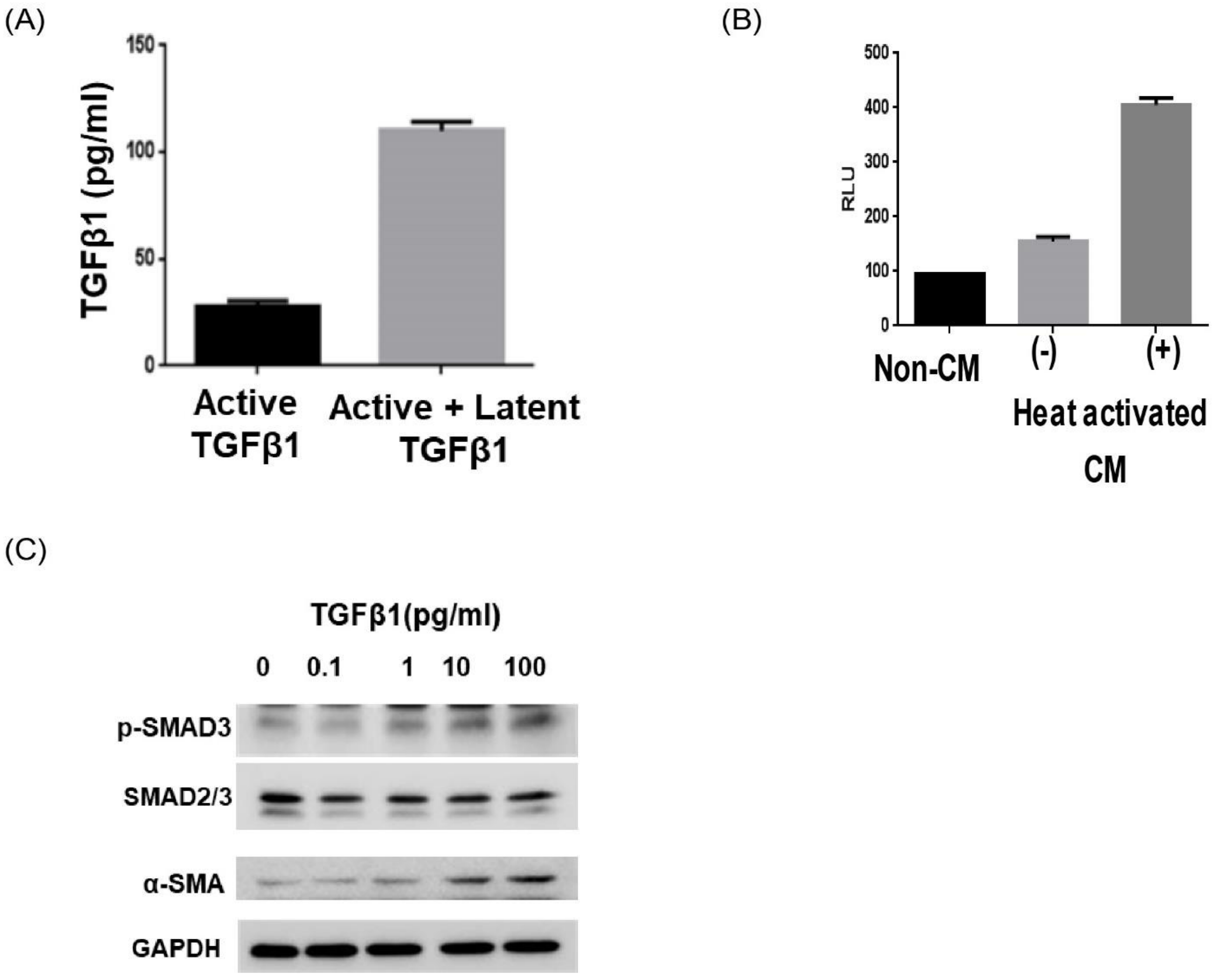

**Supplementary Figure 4. Cholesterol loading increases EVs but does not contribute to regulation of TGFβ signaling.**

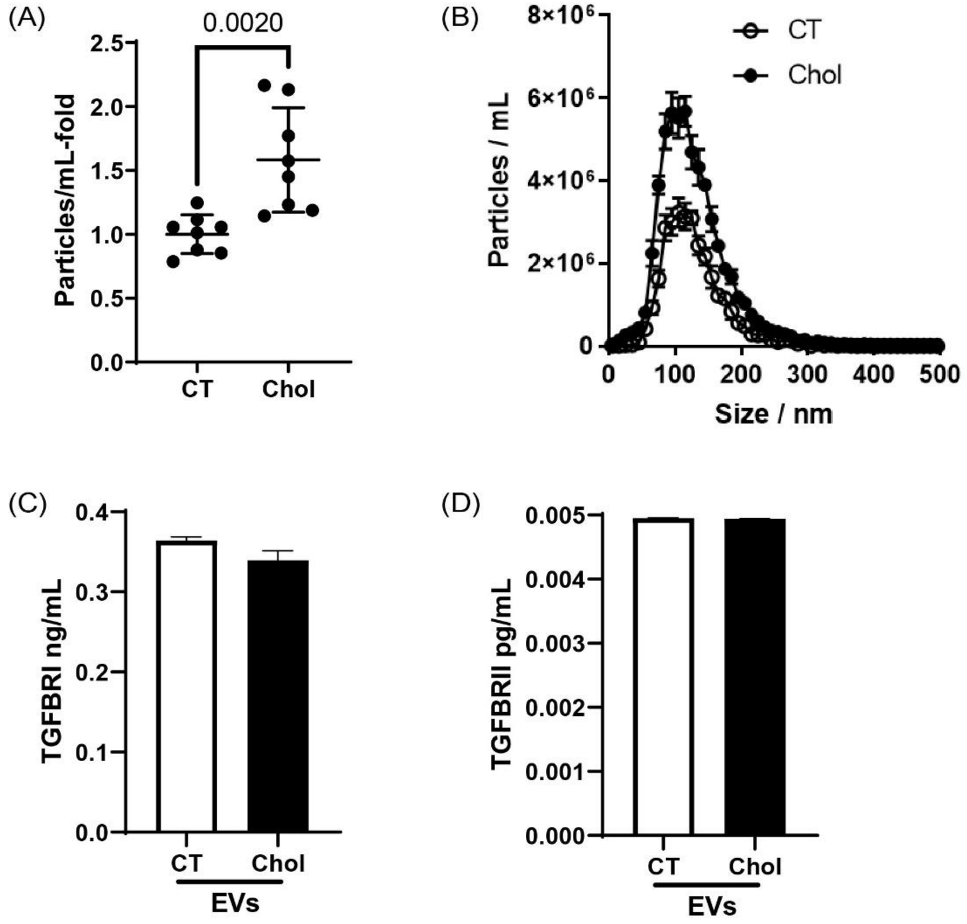

**Supplementary Figure 5. Body weight, total cholesterol and HDL cholesterol are same between TGFβR2+/+ and TGFβR2+/- mice**

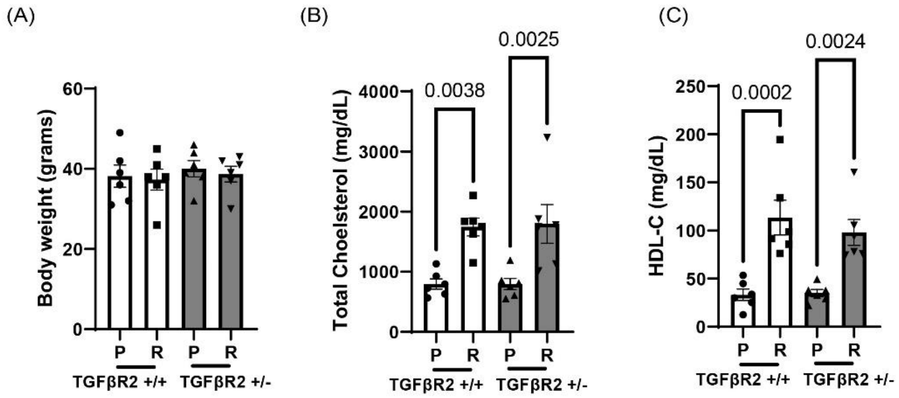

**Supplementary figure 6. HDL promotes pSMAD2 levels in TGFβR2+/- mice *in vivo***

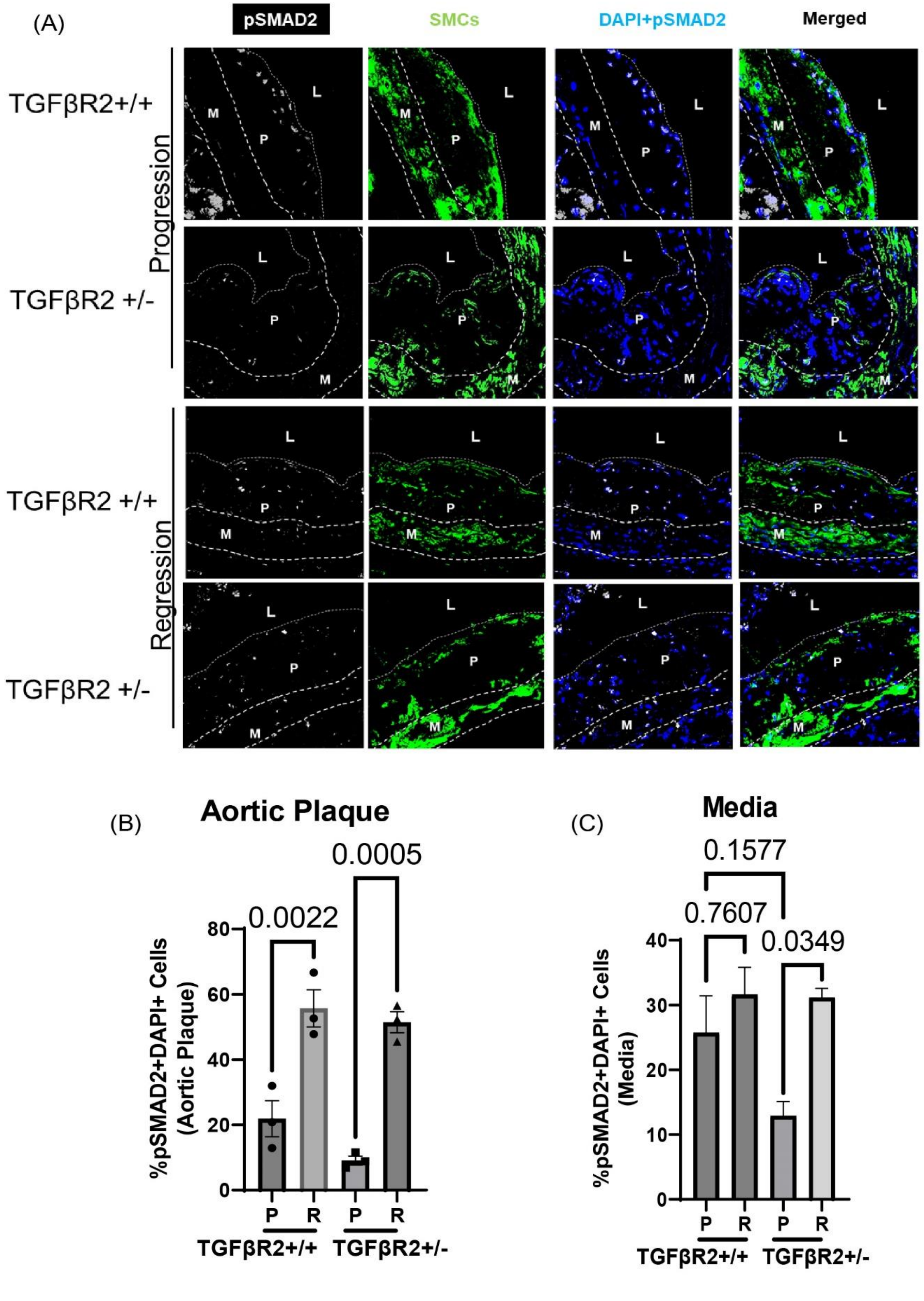

### SUPPLEMENTAL FIGURES

**S1. hVSMC loaded with cholesterol exhibited increased lipid content.** hVSMC were either untreated (Control) or treated with Cholesterol (5µg/ml) for 48h and 72h. (A) Cellular total cholesterol content was measured (B) Oil red O staining was performed to determine intracellular lipid levels and immunocytochemistry was performed to determine α-SMA expression (C) cell viability was performed using MTT assay in untreated cells (BSA) or choesltero treated cells (FC).

**S2. Cholesterol loading or using TGFβR1 antagonist did not affect the expression of total SMAD2/3 in hVSMCs.** (A) hVSMC were treated with 5µg/ml of Cholesterol for 24h or 48h. Proteins were extracted for western blotting to determine total SMAD2/3. GAPDH was used as loading control. hVSMCs were treated with (B) SB31542 (TGFβR1 antagonist) at the indicated concentration (0, 0.5, 5, 50 and 500ng/ml) for 24h. (b) Western blotting was performed to determine pSMAD2/3 levels, and total SMAD2/3. (C) RNA was extracted for real-time qPCR for *Acta2* and *Tagln*. Data are representative of two independent experiments.

**S3. hVSMCs produce active form TGFβ1.** (A) Concentration of active form or total (active + latent form) TGFβ1 was determined using ELISA (B) Conditioned medium (CM) from cultured hVSMCs were added to reporter cells which have luciferase gene driven by PAI-1 promoter in response to the stimulation of active form TGFβ1. (C) hVSMC were treated with recombinant TGFβ1 (10pg/ml) at the indicated concentration for 24h. Protein was extracted for western blotting for α-SMA. Blots was the representative of two independent experiments.

**S4. Cholesterol loading increases EVs but does not contribute to regulation of TGFβ signaling.** hVSMC were treated with Cholesterol (Chol) (5μg/ml) ot left untreated (CT) for 24h media was collected and to isolate extracellular vesicles (EVs). (A&B) Size and concentration profiles of EVs were quantified using Zetaview Nanoparticle Tracking Analysis. (C) TGFβR1 and (D) TGFβR2 expression in the isolated EVs were determined by ELISA. Data is representative of three independent experiments.

**S5. Body weight, total cholesterol and HDL cholesterol are same between TGFβR2+/+ and TGFβR2+/- mice.** Mice were placed on Western diet for 23 weeks. Before sacrificing the mice, (A) Body weight, (B) Total plasma cholesterol, and (c) Total plasma HDL-cholesterol (HDL-C) was measured (n=6 mice per group).

**S6. HDL promotes pSMAD2 levels in TGFβR2+/- mice *in vivo*.** (A) Representative image from progression (P) and regression mice (R), (TGFβR2+/+ and TGFβR2+/-) mice showing the lineage marked smooth muscle cells (GFP+ green cells) and pSMAD2 level (white color) (M-Media, P-Plauqe, L- Lumen). pSMAD2+ cells were quantified using Image J software in the (A) plaque and (B) media. Data are presented as the mean ± S.E. (n=3 mice per group).
